# Establishing And Maintaining The Blood-Brain Barrier: Epigenetic And Signaling Determinants

**DOI:** 10.1101/2022.11.29.516809

**Authors:** Jayanarayanan Sadanandan, Sithara Thomas, Iny Elizabeth Mathew, Zhen Huang, Spiros L Blackburn, Nitin Tandon, Hrishikesh Lokhande, Pierre D McCrea, Emery H. Bresnick, Pramod K Dash, Devin W McBride, Arif Harmanci, Lalit K Ahirwar, Dania Jose, Ari C Dienel, Hussein A Zeineddine, Sungha Hong, T Peeyush Kumar

## Abstract

The blood-brain barrier (BBB) controls the movement of molecules into and out of the central nervous system (CNS). Since a functional BBB forms by mouse embryonic day E15.5, we reasoned that gene cohorts expressed in CNS endothelial cells (EC) at E13.5 contribute to BBB formation. In contrast, adult gene signatures reflect BBB maintenance mechanisms. Supporting this hypothesis, transcriptomic analysis revealed distinct cohorts of EC genes involved in BBB formation and maintenance. Here, we demonstrate that epigenetic regulator’s histone deacetylase 2 (HDAC2) and polycomb repressive complex 2 (PRC2) control EC gene expression for BBB development and prevent Wnt/β-catenin (Wnt) target genes from being expressed in adult CNS ECs. Low Wnt activity during development modifies BBB genes epigenetically for the formation of functional BBB. As a Class-I HDAC inhibitor induces adult CNS ECs to regain Wnt activity and BBB genetic signatures that support BBB formation, our results inform strategies to promote BBB repair.

## Introduction

Central nervous system vessels possess a blood-brain barrier (BBB) that prevents toxins and pathogens from entering the brain. A leaky BBB can lead to deleterious consequences for CNS disorders, including stroke, traumatic brain injury, and brain tumors^1^. No treatment options are available to sustain and/or regenerate BBB integrity. Identifying and targeting the mechanism that forms and maintains the BBB is an attractive strategy to regain BBB integrity.

By comparing the EC transcriptome of peripheral and brain vessels, the genomic profile that contributes to BBB was described^2,3^. Further, extensive gene expression changes in CNS ECs during development were also reported^4,5^. However, molecular mechanisms governing BBB gene transcription and how such mechanisms contribute to BBB establishment and maintenance are unresolved.

Epigenetic modifications of histones and chromatin-modifying enzyme activities are critical determinants of gene expression^6^. Histone deacetylases (HDACs) associate with specific transcription factors and participate in gene repression^7^. HDAC inhibitors are critical experimental tools for elucidating HDAC functions^8^, and four HDAC inhibitors are FDA-approved drugs for cancer treatment^9^. More than thirty HDAC inhibitors are being investigated in clinical trials^10^. Similarly, polycomb repressive complex 2 (PRC2), a complex of polycomb-group proteins (such as EZH2, EED, and SUZ12), represses transcription via its methyltransferase activity that catalyzes tri-methylation of H3K27^11,12^. An EZH2 inhibitor is FDA-approved for Follicular lymphoma^13^.

Wnt/β-catenin (Wnt) signaling is essential for establishing the BBB^4,5,14–16^, but the activity of this pathway declines gradually after that and is reported to be minimal in adults^17–19^. Evidence linking Wnt to regulating BBB genes includes an EC-specific knockout (KO) of β-catenin that affects BBB gene expression and integrity ^5,20–22^. In CNS ECs, Wnt signaling requires binding Wnt ligands Wnt7a/7b to Frizzled receptors. This interaction stabilizes the intracellular signaling molecule β-catenin by suppressing a cytoplasmic destruction complex, which would otherwise degrade β-catenin. Stabilized β-catenin translocates to the nucleus and regulates Wnt target gene transcription by interacting with DNA binding transcription factors TCF/LEF (T-cell factor/lymphoid enhancing factor). The mechanisms underlying the reduced Wnt signaling and how Wnt regulates the BBB gene transcription are not established.

We describe the discovery of epigenetic mechanisms responsible for regulating BBB gene expression during development, how Wnt signaling is reduced, and the significance of these mechanisms for BBB development. Furthermore, we demonstrated that HDAC inhibitors activate BBB gene cohorts expressed during BBB formation, suggesting a potential therapeutic intervention.

## Results

### Transcriptional downregulation of key BBB genes in adult cortical ECs: evidence that distinct EC gene cohorts regulate BBB establishment versus maintenance

An intact, non-leaky BBB forms at E15.5^23^. We hypothesized that the EC gene cohorts involved in BBB formation are expressed on E13.5, and 3-4 month-old adult CNS EC will express gene cohorts required for BBB maintenance. We defined transcriptomes of primary cortical ECs isolated from E13.5 and 3-4-month-old adults to test this. mRNA-seq analysis revealed strong expression of EC genes, including *PECAM1*, *CDH5*, and *CLDN5,* compared to perivascular cell types such as pericytes, neuronal and glial genes (S.Fig-1A). To define the gene cohorts responsible for BBB formation vs. BBB maintenance, we identified differentially expressed genes (DEGs) expressed in natural log (fold change) in E-13.5 relative to adult mice (Fig-1A,B). DEGs with a P-value < 0.05) were considered significant. Compared to E13.5, 50% of genes were upregulated, 35% were downregulated in adults and 15% were unchanged (Fig-1C).

GO enrichment analysis of the DEGs identified five functional categories: angiogenesis, cell-to-cell junction, transporters, extracellular matrix, and DNA binding transcription factors (Fig-1D). Except for transporters, the other categories were characterized by more downregulated vs. upregulated genes. The DEGs included important BBB genes (Fig-1E). For example, tight junction (TJ) gene *CLDN1*, *CLDN5*, BBB transporters *MFSD2A*, *CAV1*, and BBB-related transcription factors *ZIC3*, *FOXF2*, and *SOX17* were differentially expressed (Fig-1E). The complete dataset is available on Geo under accession number GSE214923. Overall, the transcriptomic analysis identified EC gene cohorts that were expressed during the formation or maintenance of the BBB.

We validated the differential expression of important BBB and related genes (e.g., CLDN1, CLDN5, MFSD2A, ZIC3, and SOX17) at E-13.5, E-17.5, P0, P7, and in the adult. This analysis revealed that CLDN1, MFSD2A, and ZIC3 expressions were significantly downregulated by E-17.5 and subsequent developmental stages (Fig. 1F). Thus, the high expression of these genes might be required for the establishment and a baseline expression to maintain the BBB. By contrast, adult expression of CLDN5 was significantly upregulated compared to other developmental stages, and SOX17 was upregulated considerably compared to other developmental stages except for P7 (Fig. 1F). Additionally, we validated the mRNA expression of CLDN11 and FOXF2 (S. Fig. 1B).

**Figure 1.**
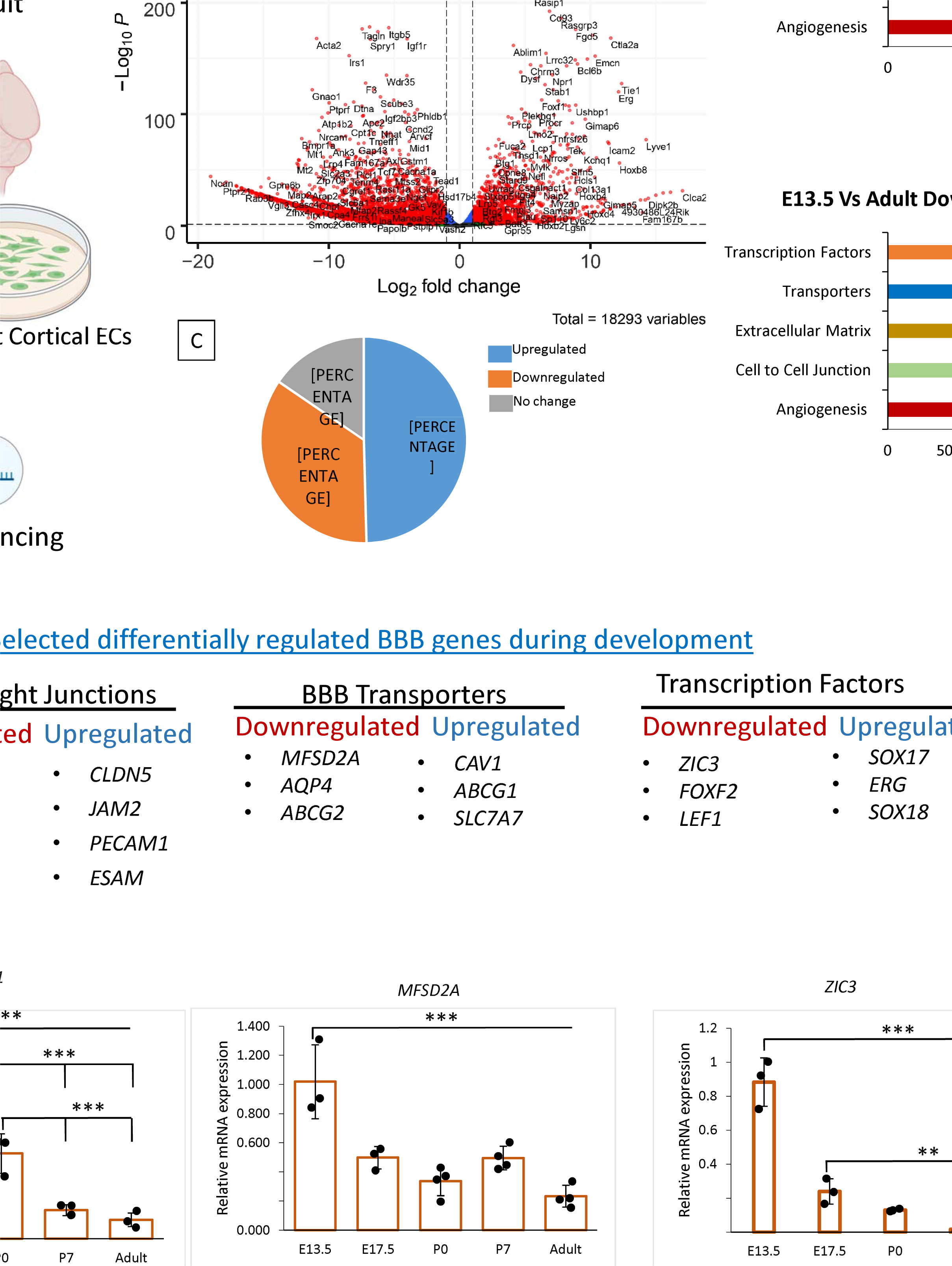
A distinct cohort of EC genes regulates the formation vs maintenance of BBB. A) Workflow for transcriptomic analysis. Primary ECs were isolated from E13.5 and adult (2-3 months old) cortex, followed by RNA isolation, mRNA sequencing, and transcriptomic analysis. B) Comparative transcriptome analysis of ECs from E13.5 and adult cortex. Volcano plot depicting downregulated and upregulated genes in adult primary cortical ECs compared to E-13.5. Genes marked in red are significant (*P*<0.05) N=3. C) Diagram depicting the percentage of genes downregulated and upregulated, and shows no differential expression in adult cortical primary ECs compared to E-13.5. D) Downregulated and upregulated genes were categorized with five important EC functions. E) Enrichment analysis revealed important BBB-related genes that were differentially regulated during development. F) Relative mRNA expression of *CLDN1*, *MFSD2A, ZIC3*, *SOX17,* and *CLDN5*. in primary cortical ECs isolated from E13.5, E-17.5, P0, P7 and adult. Significant differences are observed between E-13.5 and all consecutive stages for *CLDN1*(*** *p*<0.001, N=3/group), *MFSD2A* (*** *p*<0.001, N=3-4/group) and ZIC3(*** *p*<0.001N=3-4/group). CLDN1 showed significant differences between E-17.5 vs P7 and adult and P0 vs P7 and adult (*** *p*<0.001). ZIC3 significantly differed between E-17.5 vs P7 and adult es (** *p*<0.01). Significant differences are observed between E-13.5 vs P7 and adults for *CLDN5* and *SOX17* (* *p*<0.001 N=3-4/group).

Hupe *et al* ., (2017) demonstrated differentially regulated genes in brain ECs during embryonic development^5^. To further validate our data, we compared our transcriptome with the 1129 genes shown to be regulated during embryonic development. We found 315 of our downregulated transcripts in adults and 490 of our upregulated transcripts in adults matched with their dataset. Key overlapping genes are highlighted (S. Fig. 1C).

### HDAC2 and PRC2 mediated transcriptional regulation of BBB genes

Histone deacetylases (HDACs) have critical roles in development and tissue homeostasis, and HDAC inhibitors are instructive experimental tools^24,25^. To assess whether the transcriptional downregulation of BBB genes involves HDAC-dependent epigenetic repression, we treated adult ECs with a pan HDAC inhibitor trichostatin A (TSA) for 48 hours. TSA increased *CLDN1* (Fig-2A) and *ZIC3* (S.Fig-2A) expression relative to the control. Since there are four major HDAC classes, class I (HDAC 1, 2, 3, and 8), class II (HDAC 4, 5, 6, 7, 9, 10), class III (SIRTs 1–7), and class IV (HDAC 11), we tested whether specific HDACs mediate the repression. The class-I HDAC inhibitor MS-275 significantly upregulated *CLDN1*, MFSD2A and ZIC3 while downregulated *CLDN5* (Fig-2A, S.Fig-2A). No significant difference in CLDN1 mRNA expression was observed with class-II HDAC inhibitor (S.Fig-2A). In adult ECs, HDAC2 exhibited greater expression than other class I HDACs (Fig-2B). To analyze the HDAC2 function, we utilized siRNAs to downregulate HDAC1, HDAC2, or HDAC3 in adult ECs. Knockdown (KD) of HDAC2 significantly upregulated the repressed *CLDN1*, *MFSD2A*, and *ZIC3* genes and reduced *CLDN5* expression (Fig-2C). By contrast, HDAC1 and HDAC3 downregulation did not affect these genes (not shown). BBB genes analyzed above were selected based on their expression patterns and to represent important functional attributes of BBBs, such as tight junctions (*CLDN1* and *CLDN5*), transporters (*MFSD2A*), and transcription factors (*ZIC3*). We used ChIP-qPCR to test whether HDAC2 directly regulates BBB gene expression in E13.5 and adult cortical EC contexts. Three primers were designed ((-)500, TSS & (+)500) spanning the 1 kb region on each side of the promoter. HDAC2 occupancy was detected in the (-)and(+) 500 regions of *CLDN1* in adults with no significant enrichment in E13.5. *ZIC3* showed occupancy at all three regions (1kb) in the adult stage, with E13.5 showing enrichment in TSS only (Fig-2D, G &S.Fig2B). Conversely, HDAC2 occupied *MFSD2A* only at E13.5((-)500 & TSS), and in *CLDN5* occupancy was detected at both E13.5 (1kb) and in adults ((-) & (+) 500). These results link HDAC2 to the developmental control of BBB genes (Fig-2D,G & S.Fig2B).

Our initial screening showed that the PRC2 inhibitor DZNEP significantly increased *CLDN1* expression in adult ECs (S.Fig-2A). To analyze the PRC2 function in this context, we downregulated the PRC2 subunit EZH2 from adult ECs. Compared to the control, EZH2 downregulation significantly increased *CLDN1*, *ZIC3*, and *MFSD2A* expression and decreased *CLDN5* expression (Fig 2E). ChiP-qPCR of PRC2 subunit EED revealed EED occupancy at various regions of *CLDN1*, *MFSD2A,* and *ZIC3* at E13.5 and in the adult (Fig2F,G & S.Fig-2). EED occupied *CLDN5* at E13.5, but not in adult ECs. Our data support a model in which HDAC2 and PRC2 are critical determinants of EC BBB transcriptomes during BBB development.

**Figure 2.**
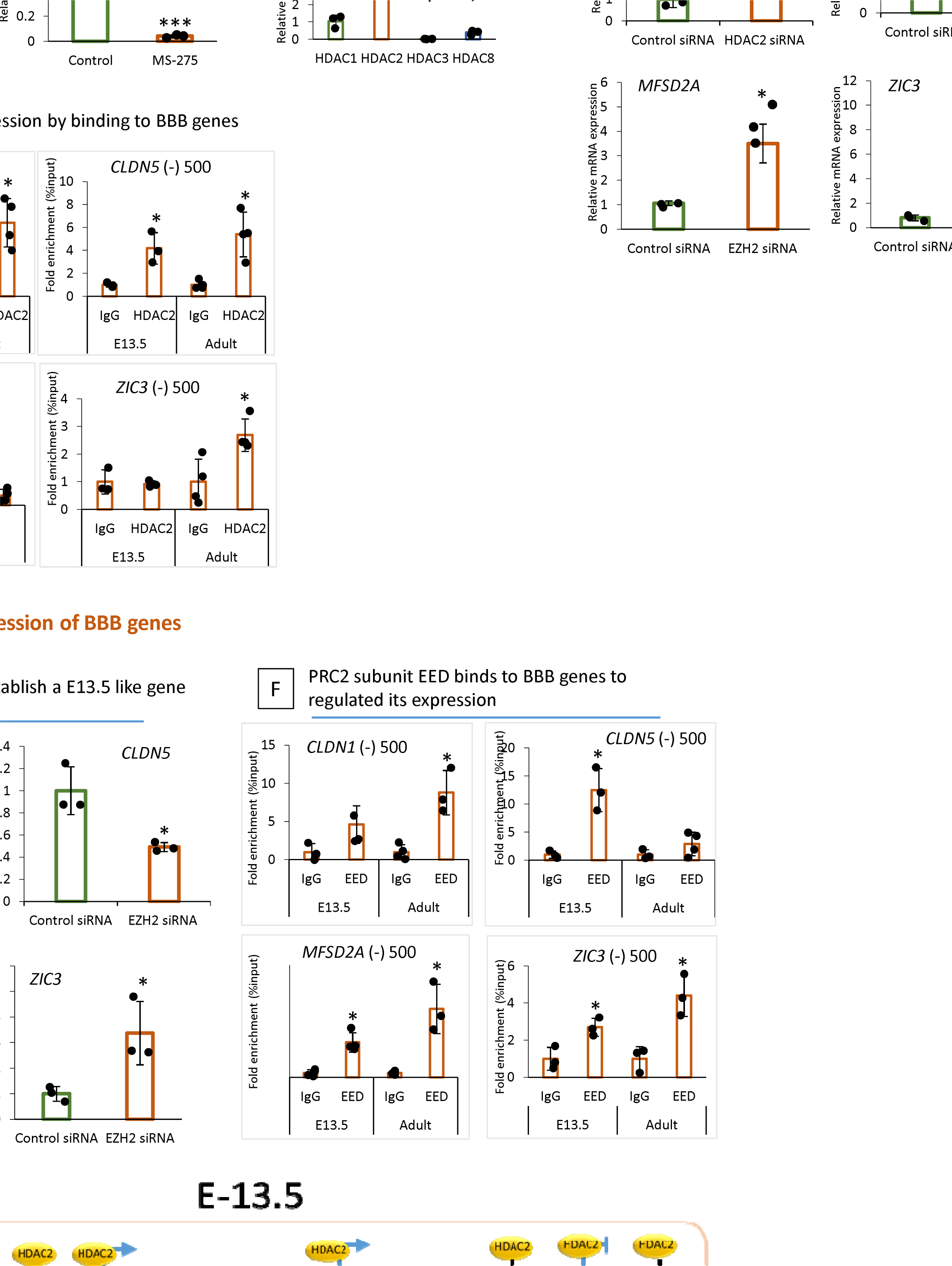
Epigenetic regulators HDAC2 and PRC2 regulate the transcription of BBB genes. A) qPCR of adult primary cortical ECs treated with TSA (200nm) and MS-275(10um) 48hrs showed significantly increased mRNA expression of *CLDN1* compared to DMSO treated control while *CLDN5* was significantly decreased with MS-275 treatment when compared to control (* *p*<0.001 vs Control #*p*<0.001 vs TSA treatment N=3/group). B) mRNA expression level of class-I HDAC family members in adult primary cortical ECs. Expression was normalized to housekeeping genes GAPDH and HDAC1. Significant mRNA expression of HDAC2 was observed in adult primary cortical ECs compared to other Class-I HDACs (* *p*<0.05 vs HDAC1,3 and 8 N=3/group). HDAC1 showed significantly higher expression compared to HDAC3 and HDAC8 (* *p*<0.05) and HDAC8 showed significantly higher expression compared to HDAC3 (* *p*<0.05). C) Effect of HDAC2 siRNA on BBB gene expression in adult cortical ECs. Using lipofectamine adult cortical ECs were transfected with HDAC2 siRNA (500µg). qPCR analysis revealed that compared to control siRNA treated group HDAC2 siRNA treated ECs showed significantly increased expression of *CLDN1*(* *p*<0.001), *MFSD2A* (* *p*<0.05) and *ZIC3* (* *p*<0.05) while *CLDN5* (* *p*<0.05) showed significantly decreased expression. N=3/group D) HDAC2 occupancy of the indicated chromatin regions in primary cortical ECs from E-13.5 and adult. Occupancy was measured by ChIP followed by quantitative PCR (ChIP-qPCR). The adjacent gene and the distance to the TSS name chromatin regions. (* *p*<0.05 vs IgG N=3-4/group) E) qPCR analysis of *CLDN1*, *CLDN5*, *MFSD2A,* and ZIC3 in EZH2 and control siRNA-treated adult primary cortical ECs. Compared to the control siRNA-treated group, EZH2 siRNA-treated ECs showed significantly increased expression of *CLDN1* (* *p*<0.001), *MFSD2A* (* *p*<0.001), and *ZIC3* (* *p*<0.05), while *CLDN5* (* *p*<0.05) showed significantly decreased expression. N=3/group. F) ChIP-qPCR analysis of PRC2 subunit EED on indicated chromatin regions and genes in E13.5 and adult primary cortical ECs. * *p*<0.05 vs IgG, N=3-4/ group. G) Schematic representation of HDAC2 and PRC2 binding on indicated genes in cortical ECs at E-13.5 and adult.

**Figure 3.**
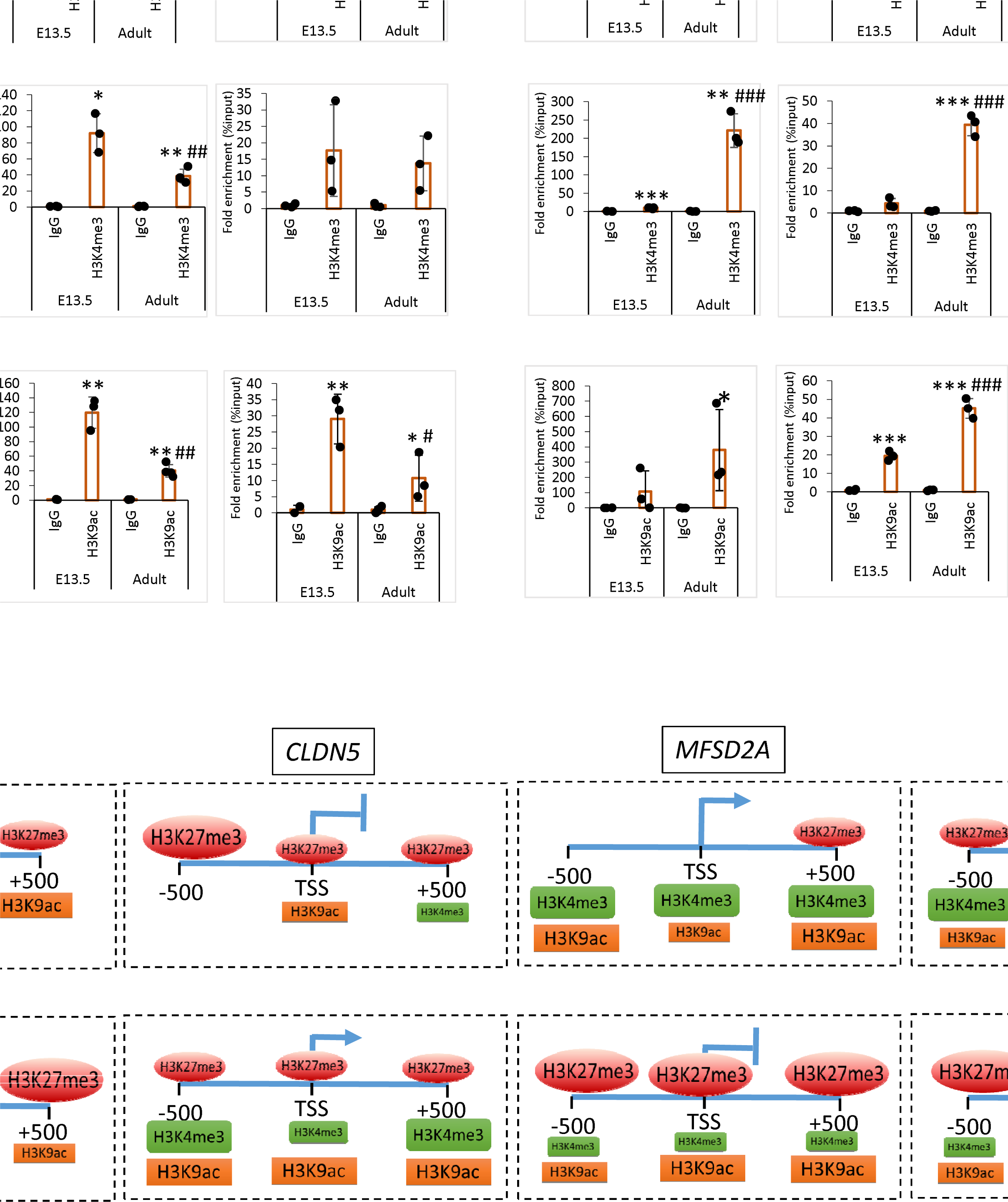
BBB genes exhibit diverse post-translational histone modifications. A) ChIP-qPCR analysis of histone marks H3K27me3, H3K4me3, and H3K9ac on 1 KB region *CLDN1* gene in E13.5 and adult cortical ECs. *CLDN1* gene was downregulated in ECs during development. The ChIP signals were normalized to IgG. B) ChIP-qPCR showing the H3K27me3, H3K4me3, and H3K9ac density at the CLDN5 gene in E13.5 and adult cortical ECs. CLDN5 gene was upregulated during development. Data are shown as mean ± S.D. \*\*\**p* < 0.001, \*\**p* < 0.01, \**p* < 0.05 vs IgG & ###*p* < 0.001, ##*p* < 0.01, #*p* < 0.05 vs respective E-13.5 histone mark. N = 3-4/group. C) Schematic representation of H3K27me3, H3K4me3, and H3K9ac binding density on *CLDN1*, *CLDN5*, *MFSD2A*, and *ZIC3* in E13.5 and adult primary cortical E.C.s. Shape size indicates the binding density in the indicated chromatin regions.

### Distinct histone modifications delineate the transcription program of BBB genes

Since HDAC2 and PRC2 regulate multiple BBB genes, we sought to determine whether they share similar post-translational histone modifications. To examine this, we performed ChIP-qPCR of potentially involved histone marks, including the repressive marks H3K27me3, H3K9me3, and the activating marks H3K4me3 and H3K9ac. We have scanned approximately 1kb genomic region surrounding the TSS of *CLDN1*, *CLDN5*, *MFSD2A*, *ZIC3,* and *SOX17* in E13.5 and adults. H3K9me3 ChIP-qPCR on our selected BBB genes didn’t show significant binding in either stage (S.Fig-3A). We detected abundant enrichment of repressive histone mark H3K27me3 on *CLDN1* (Fig-3A,C), *MFSD2A*, and *ZIC3* (Fig-3C & S.Fig-3B, C) in adult E.C.s compared to E13.5. *CLDN1, MFSD2A,* and *ZIC*3 showed significant enrichment of H3K27me3 compared to IgG in E13.5 (Fig-3A & S.Fig-3B,C). While *CLDN5* showed significant enrichment of H3K27me3 in the 1kb region in both E13.5 and adults (Fig-3B). However, E13.5 showed abundant enrichment of H3K27me3 in the -500 region compared to adults (Fig-3B).

Active histone mark, H3K4me3 showed significantly increased enrichment on *CLDN1* (TSS), *MFSD2A* (1 KB), and *ZIC3* (-500 and +500) at E13.5 compared to adult (Fig-3A and S.Fig-3B, C). *CLDN1* (-)500 region didn’t show any significant binding for H3K4me3 in both stages, while the (+)500 region showed significant binding compared to IgG with no difference between E-13.5 and adult. *ZIC3* TSS region showed significant enrichment of H3K4me3 in the adult compared to E-13.5(S.Fig-3). Supporting its abundant expression in adults, *CLDN5* (1 KB) showed significant enrichment of H3K4me3 in adults compared to E13.5. Another active histone mark, H3K9ac, showed significant enrichment on *CLDN1* (TSS and +500) at E13.5 compared to the adult with no significant binding in the -500 region in both stages (Fig-3A & S.Fig-3). *ZIC3* (TSS and +500) showed a significant binding for H3K9ac in E13.5 compared to the adult, while the -500 region didn’t show any enrichment in both stages. *MFSD2A* showed significant enrichment of H3K9ac in the -500 and +500 regions at E-13.5 compared to adults. At the same time, the TSS region showed significant enrichment in adults compared to E-13.5. Histone modifications on *SOX17* are shown in S.Fig-3D. These results indicate that BBB genes acquire a unique epigenetic signature during development.

### HDAC2 activity is critical for the maturation of BBB, while PRC2 is dispensable

To examine the role of HDAC2 and PRC2 in BBB maturation, we knock out (KO.) *HDAC2* and PRC2 subunit *EZH2* from ECs during embryonic development. To conditionally KO *HDAC2* or *EZH2,* we used tamoxifen-inducible Cdh5(PAC)-CreERT2 mice. Tamoxifen was injected into the mother starting at E-12.5 and on alternate days until E-16.5 (Fig-4A & 4E). This allowed the KO of *HDAC2* or *EZH2* before the maturation of the BBB.

It was observed that *HDAC2 and EZH2* KO embryos grew normally and were alive on the day of sacrifice E17.5. Significantly decreased mRNA expression of *HDAC2* in the whole brain analysis confirmed the loss of HDAC2 (S.Fig-4). Compared to WT, *HDAC2* ECKO pial vessels were dilated and showed increased angiogenesis (Fig-4B). BBB permeability was assessed in *HDAC2* ECKO mice using a 70KD FITC-conjugated dextran tracer. The tracer remained confined inside the vessels of E17.5 WT embryos, supporting previous findings that the BBB matures by E-15.5 (Fig-4C). Conversely, in HDAC2 ECKO, dextran leaked into the cortical parenchyma (Fig-4C). Green fluorescent intensity measurements confirmed the immature BBB in the *HDAC2 ECKO* (Fig-4D). A vascular analysis using angiotool determined that HDAC2 ECKO had a significantly higher vascularized brain area than WT, indicating that HDAC2 plays a key role in angiogenesis (Fig-4D). The pharmacological inhibition of class I HDAC using MS-275 in timed pregnant WT mice at E-13.5 showed significant tracer leakage into the brain parenchyma at E-15.5 compared to vehicle-treated control (S.Fig-4A). The MS-275 treated embryos also showed a thin cortex compared to the control, possibly due to the leakage of the drug into the brain which affects brain development (S.Fig-4A). These findings should be considered when considering the clinical use of MS-275 in a developing brain.

Compared to WT, at E-17.5 *EZH2 ECKO* showed dilated vessels with no evident increase in the pial vessel angiogenesis (Fig-4F). Confirming the loss of *EZH2*, mRNA analysis on the whole brain showed a significant reduction. BBB permeability assay showed a subtle leakage of 70KD fluorescent tracer into the brain parenchyma (Fig-4G). However, the fluorescence intensity analysis showed no significant difference (Fig-4H). Further, the mRNA analysis on the whole brain showed a significant decrease in *EZH2* expression with no difference in *CLDN1*, *MFSD2A*, and *CLDN5* (S.Fig-4C). Together, *HDAC2* and *EZH2* ECKO data suggest that HDAC2 is critical for BBB maturation, whereas PRC2 is dispensable or a support mechanism required during later BBB development.

### Despite Wnt pathway activity in the adult, Wnt target genes are epigenetically repressed

Wnt pathway has been shown to influence BBB gene expression^5^. However, this pathway activity is reported to be minimal in adults^15,18^. In our transcriptomic analysis, 67% of Wnt signaling-related genes were downregulated in adults, while 33% were upregulated (Fig-5A). Interestingly, downstream Wnt targets genes including *AXIN2*, *LEF1*, and *VEGFA* have downregulated in adult ECs, while the upstream Wnt pathway components like *FZD4/6*, *LRP5*, and *CTNNB1* were upregulated (Fig-5a). Since Wnt-regulated BBB genes such as *CLDN1*, and *ZIC3* expression was also minimum in adults, we hypothesize that the Wnt pathway is still active in adult CNS ECs. In contrast, Wnt target genes are epigenetically repressed.

To test this, we activated the Wnt pathway in E13.5 and adult ECs using identical concentrations of Wnt3a ligand or GSK3B inhibitor CHIR99021. mRNA analysis revealed that Wnt target genes *AXIN2* and *LEF1* can be significantly activated in E13.5 ECs when treated with Wnt3a or CHIR99021, while no significant activation was observed in adults (Fig-5B). Further validating this finding, transcriptome analysis on adult ECs after Wnt3a treatment showed only activation of one Wnt target gene CD44 (Fig-5C). Intriguingly, the Wnt3a treatment downregulated 16 Wnt-related genes (Fig-5C). Next using immunohistochemistry of β-catenin in adult control and CHIR 99021 treated ECs we demonstrated that Wnt pathway activation could translocate or stabilize the Wnt transducer β-catenin into the nucleus (Fig-5D). These results indicate that the Wnt pathway is active in adult CNS ECs.

**Figure 4.**
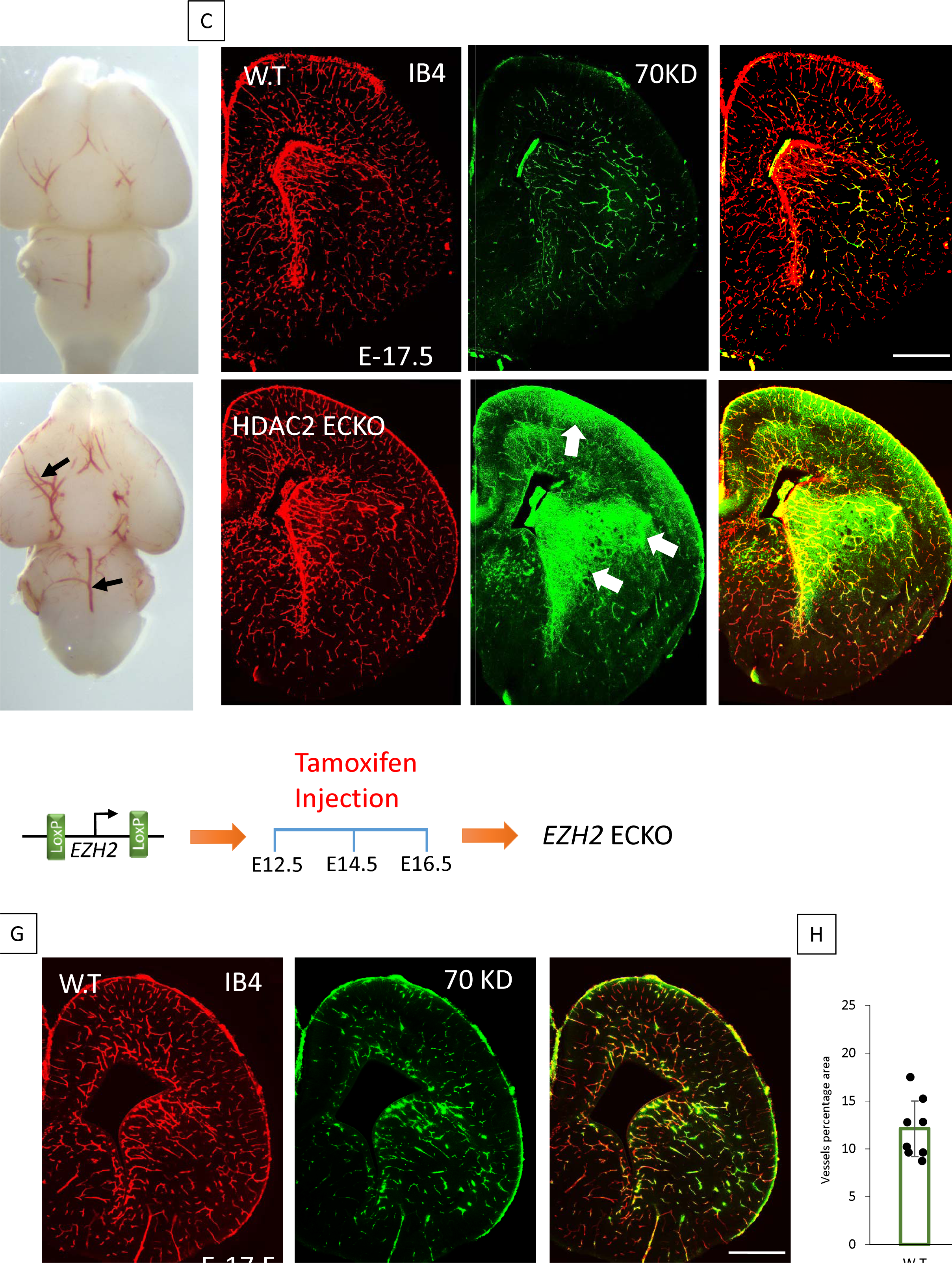
HDAC2 activity is required to form a functional BBB, while PRC2 is dispensable. A) Effect of deletion of HDAC2 from the ECs during embryonic development. Schematic representation of breeding scheme and generation of EC-specific KO of HDAC2. Tamoxifen was delivered to the pregnant mother at E-12.5, and brains were harvested at E-17.5. B) Representative phase microscopy images of the dorsal surface and ventral surface of the brain at E-17.5. HDAC2 ECKO shows a significant increase in pia vessels, as pointed out by black arrows on the dorsal surface. Black arrows in the ventral surface showed dilated vessels. C) BBB permeability assay using 70KD tracer and isolectin B4 (IB4) staining to image the vessels. In HDAC2 ECKO, a green fluorescent tracer leaked out of the vessels, as indicated by the white arrows. 10x images are acquired and merged using tile scanning. Scale bar, 500µm D) Vessel percentage area of HDAC2 ECKO was significantly higher than WT (* *p*<0.001 vs control N=10). A significant increase in fluorescent intensity was quantified in HDAC2 ECKO compared to WT (* *p*<0.0001 vs control N=3 for WT and N =5 for HDAC2 ECKO). E) Schematic representation of breeding scheme and generation of EZH2 ECKO. Tamoxifen was injected the same as for HDAC2 ECKO. F) Representative phase microscopic image of WT and EZH2 ECKO. As black arrows show, the EZH2 ECKO brain shows dilated vessels with no visible increase in pial angiogenesis. G) BBB permeability assay using 70KD-FITC Dextran shows subtle tracer leakage out of the vessels in EZH2 ECKO, represented by white arrows. IB4 staining reveals the vessels in the brain. 10x images are acquired and merged using tile scanning. Scale bar, 500 µm H) Quantification of vessel percentage area (WT N=9 EZH2 ECKO N=10) and fluorescent intensity didn’t show any significant difference between WT and EZH2 ECKO (WT N=5 EZH2 ECKO N=5).

Next, we investigated whether Wnt target genes are epigenetically repressed. Wnt target genes *AXIN2* and *LEF1* were significantly upregulated in adult CNS ECs when treated with HDAC2 or EZH2 siRNA, MS-275, and in HDAC2 ECKO mutants (Fig-5E & S.F-5A,B). Furthermore, the *AXIN2* promoter in adults showed significantly increased occupancy of HDAC2, repressive histone mark H3K27me3 in adults while active histone marks H3K4me3 and H3K9ac were reduced considerably compared to E-13.5 as analyzed by ChIP-qPCR (Fig 5F). *LEF1* also showed a similar repressive histone modification (Fig-5C). Further, MS-275 and LiCl (Wnt agonist) treatment increase the AXIN2 protein expression in adult cortical vessels but not by just LiCl (S.Fig-5D). Thus, our data explain the mechanism behind the low Wnt pathway in adult CNS ECs.

**Figure 5.**
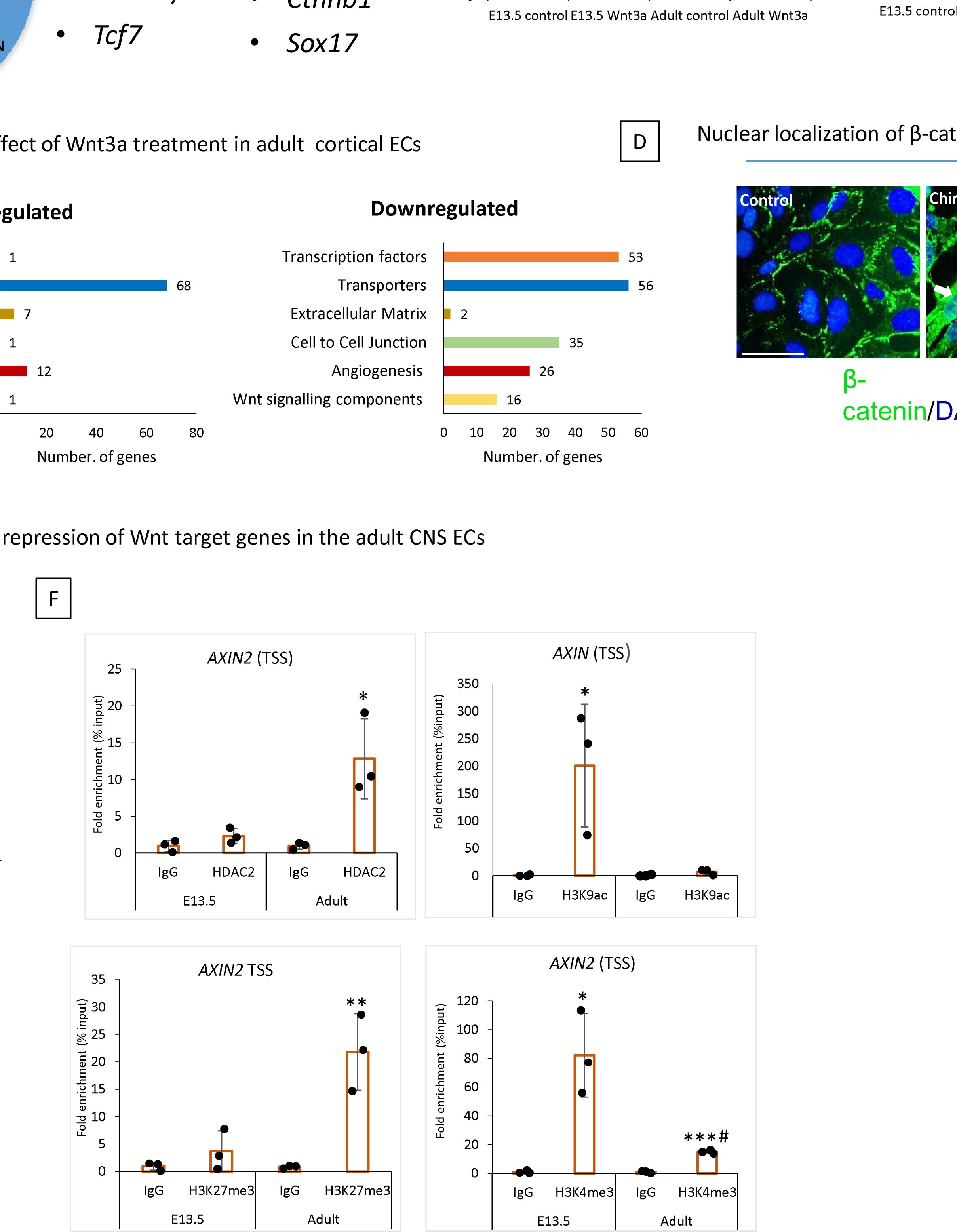
Wnt pathway is active in adult CNS ECs, but the Wnt target genes are epigenetically repressed. A) Diagram depicting the percentage of Wnt-related genes downregulated and upregulated adult cortical primary ECs compared to E-13.5. Selected important Wnt-related genes are shown. B) Ligand-independent transcriptional repression of Wnt target genes. In primary cortical ECs from E-13.5, activating the Wnt pathway with Wnt3a (200ng/mL) or CHIR- 99021 (5uM) for 48 hrs. caused increased mRNA expression of Wnt target genes *AXIN2* and *LEF1* (measured via qRT-PCR). However, activation of the Wnt pathway in primary adult mouse brain ECs does not increase Wnt target gene expressions. *p<0.001 vs E-13.5 control, ns-no significant difference n=3/group. C) mRNA sequencing was performed in control and Wnt3a (200ng/mL) treated adult primary cortical ECs. Differentially expressed genes were categorized into six categories important to CNS endothelial cells. D) Immunofluorescence staining of β- catenin (green) in control and Wnt agonist Chir-99021 treated endothelial cells. White arrows indicate the nuclear localization of β-catenin to the nucleus. 20x images are acquired, cropped, and enlarged. Scale bar, 1 µm. E) Adult primary cortical ECs transfected with control, HDAC2 & EZH2 siRNA showed significant upregulation of Wnt target genes AXIN2. *\*\*\*p*<0.001 vs control siRNA & *#p*<0.05 vs control siRNA. N=3/group F) HDAC2, histone marks H3K27me3, H3K4me3, and H3K9ac occupancy on the AXIN2 TSS regions in primary cortical ECs from E-13.5 and adult. ChIP-qPCR measured occupancy. N=3/group \*\*\**p* < 0.001, \*\**p* < 0.01, \**p* < 0.05 vs IgG & #*p* < 0.05 vs H3K4me3 E-13.5.

**Figure 6.**
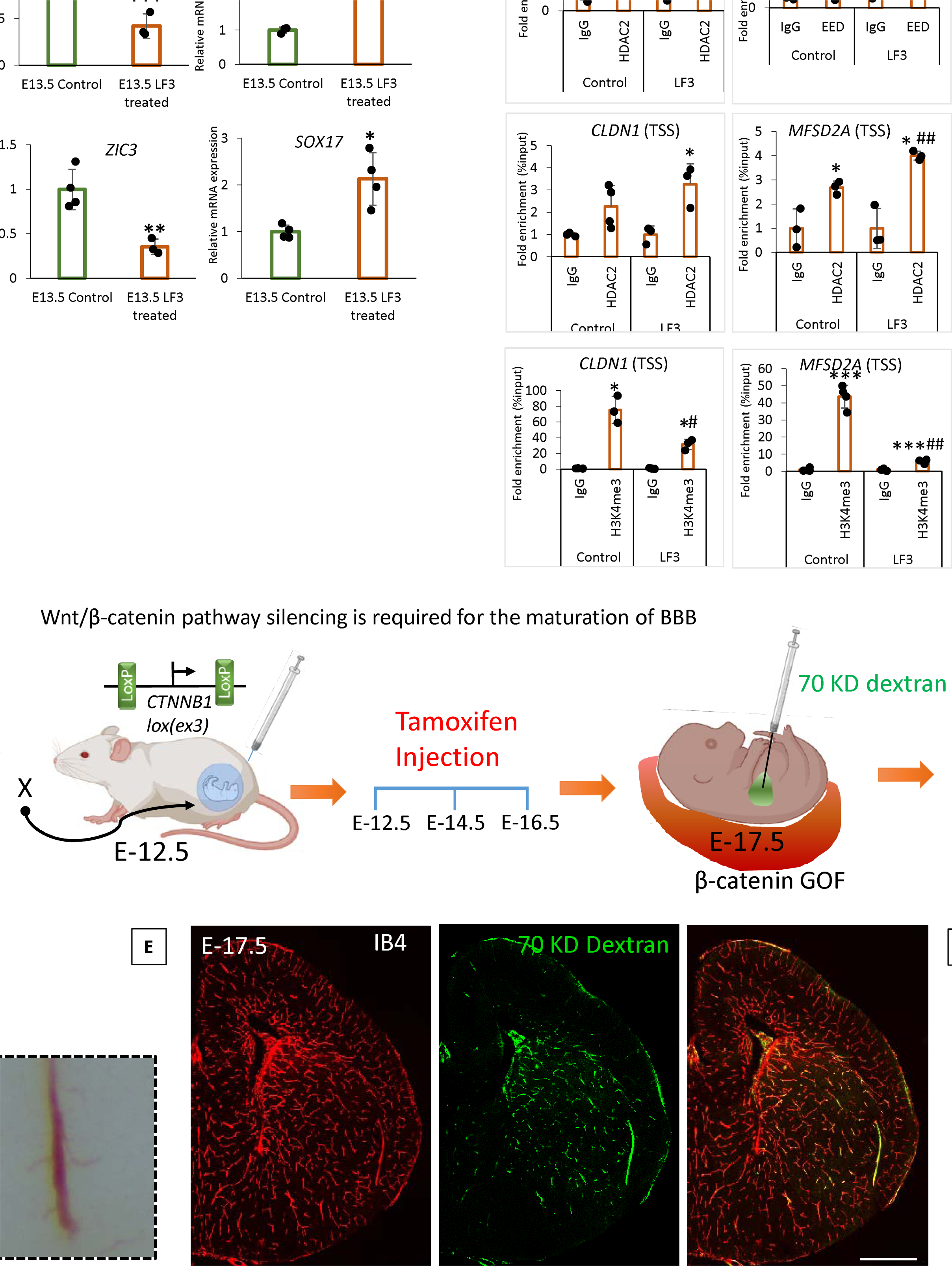
Low Wnt signaling epigenetically modifies the BBB genes to achieve BBB maturation. A) Effect of Wnt pathway inhibition on BBB genes in E13.5 primary cortical ECs. E-13.5 ECs were treated with LF3(50um) for 48 hours to inhibit the Wnt pathway. Significantly decreased mRNA expression of *AXIN2* confirmed the reduced Wnt pathway. mRNA expression of *CLDN1*, *MFSD2A* and *ZIC3* was significantly decreased *CLDN5* and *SOX17* expression was significantly increased after LF3 treatment. Data are shown as mean ± S.D. \*\*\**p* < 0.001, \*\**p* < 0.01, \**p* < 0.05 vs E13.5 control N=3/group. B) Wnt pathway inhibition *via* LF3 induce epigenetic modifications in target gene *AXIN2* and BBB genes *CLDN1*, *MFSD2A*, and CLDN5. First row- *AXIN2* showed significant enrichment of HDAC2 and EED in LF3 treated E13.5 ECs compared to control (\**p* < 0.05 vs IgG). Histone mark H3K27me3 showed significant enrichment in both conditions compared to IgG however, LF3 treated ECs showed significantly increased enrichment compared to the control. \*\**p* < 0.01 vs IgG, #*p* < 0.05 vs control N=3-4/group. Second row- LF3 treated E13.5 ECs showed significant enrichment of HDAC2 in *CLDN1* TSS (\**p* < 0.05 vs IgG N=3-4/group). *MFSD2A* showed significant enrichment in both conditions, while LF3 treatment showed a significantly increased enrichment compared to the control (\**p* < 0.05 vs IgG & ##*p* < 0.01 vs control N=3/group). *CLDN5* didn’t show any significant difference in HDAC2 binding (not shown). At the same time, active histone marks H3K9ac showed significant enrichment in both conditions with an increased enrichment with LF3 treatment (\*\*\**p* < 0.001, \**p* < 0.05 vs IgG & #*p* < 0.05 vs control N=3/group). Third row- H3K4me3 ChIP-qPCR on the TSS region of *CDLN1*, *MFSD2A,* and *CLDN5* showed significant enrichment in both conditions with a decreased enrichment with LF3 treatment on *CDLN1* and *MFSD2A* and an increased enrichment with LF3 treatment on *CLDN5*. \*\*\**p* < 0.001, \**p* < 0.05 vs IgG & ##*p* < 0.01, #*p* < 0.05 vs control N=3-4/group. C) Schematic representation of breeding scheme and generation of EC-specific gain of function (GOF) of β-catenin. Tamoxifen was delivered to the pregnant mother at E-12.5 and brains were harvested at E-17.5. D) Representative Phase microscopy image of the dorsal brain from WT and β-catenin-GOF. Images in the square box were enlarged to show increased pial vessel angiogenesis. E) BBB permeability assay using the 70KD FITC-Dextran tracer. Cortical vessels were stained using IB4.10X tile scanning images acquired and merged. FITC dextran was leaked out of the vessels in the brain of β-catenin GOF compared to WT 10x images were acquired and merged using tile scanning. Scale bar, 500 µm F) Quantification of brain vessels percentage area (\*\*\**p* < 0.001 vs WT N=4-5/group) and green fluorescent intensity showed a significant increase in β-catenin GOF compared to WT \*\*\**p* < 0.001, \**p* < 0.05 vs WT N=5- 6/group.

### Low Wnt signaling epigenetically modifies the BBB genes to achieve BBB maturation

We investigated the relevance of the low Wnt pathway to BBB development. To this, we treated primary E13.5 cortical ECs with LF3 (inhibits the interaction of β-catenin and TCF4) for 48 hours. LF3 activity was confirmed by reduced mRNA expression of the Wnt target gene *AXIN2* (Fig-6A). LF3 treatment induces adult/BBB maintenance-type gene expression patterns in E13.5 with a significant decrease in *CLDN1*, *ZIC3*, and *MFSD2A* expression and an increase in *CLDN5* and *SOX17* expression (Fig-6A). Another Wnt inhibitor IWR-1-endo also showed similar results (not shown), while β-catenin siRNA KD showed a similar gene expression pattern for AXIN2, CLDN1, ZIC3, and MFSD2A and no difference in *CLDN5* (S.Fig-6A).

To determine whether Wnt pathway inhibition induces epigenetic modifications on its target genes and BBB genes, we performed Chip-qPCR analysis of HDAC2, EED, and histone mark H3K27me3, H3K4me3, and H3K9ac on the promoter of Wnt target genes *AXIN2*, and *LEF1*, BBB genes *CLDN1*, *CLDN5*, *MFSD2A,* and *ZIC3*. An increased HDAC2 occupancy was observed on AXIN2, LEF1, CLDN1, and MFSD2A promoter when E13.5 ECs were treated with LF3 (Fig-6B & S.Fig-6B). *ZIC3* and *CLDN5* showed no significant difference (not shown). EED and histone mark H3K27me3 showed an increased enrichment on the *AXIN2 while LEF1* showed increased enrichment for H3K27me3 in the promoter after LF3 treatment (Fig-6B & S.Fig-6B) with no difference in other genes analyzed (not shown). *CLDN1* and *MFSD2A* showed a significant decrease in the enrichment of active histone mark H3K4me3 after treatment with LF3, whereas *CLDN5* showed a significant increase (Fig-6B). Another active histone mark H3K9ac showed significantly increased enrichment on *CLDN5* (Fig-6B) with no difference in the promoter of other genes analyzed (not shown).

We then investigate the significance of the physiological reduction of Wnt signaling on BBB maturation. For this we use Ctnnb1lox(ex3)^+/+^; Cdh5-CreERT2 mice which are widely used to attain the inducible EC specific β-catenin gain of function (GOF) (Fig-6C). Upon tamoxifen treatment exon 3 of *CTNNB1* (encoding β-catenin) will be deleted, leading to the expression of a stabilized form of β-catenin protein and thereby constitutive activation of canonical Wnt signaling (Fig6C). Tamoxifen was injected into the pregnant mother, as explained in section 2.4. At E-17.5, β-catenin GOF embryos showed significantly increased pial angiogenesis (Fig-6D) compared to WT BBB permeability assay using 70KD FITC dextran revealed immature BBB in β-catenin GOF (Fig-6E & F). Confirming this result, the pharmacological activation of Wnt signaling by the Wnt agonist LiCl also showed a significant BBB leakage compared to the control (S.Fig- 6C). Our data suggest that a low Wnt pathway supports BBB maturations by epigenetically modifying BBB genes.

Embryonic deletion of EC HDAC2 and Class-I HDAC inhibitor treatment of adult CNS ECs activate gene expression related to angiogenesis, BBB formation, and the Wnt pathway.

Since the E-17.5 HDAC2 ECKO showed a significant increase in vessel density and BBB leakage, we hypothesize this is due to the inefficient epigenetic repression of angiogenesis and BBB formation genes during development. To investigate this, we utilized E-17.5 HDAC2 ECKO and FACS-sorted CD31+ ECs for ultra-low mRNA sequencing (Fig. 7A). Indicating the critical role of HDAC2 in regulating the EC gene expression 2257 genes show significant upregulation and 1723 genes showed downregulation.

**Figure 7.**
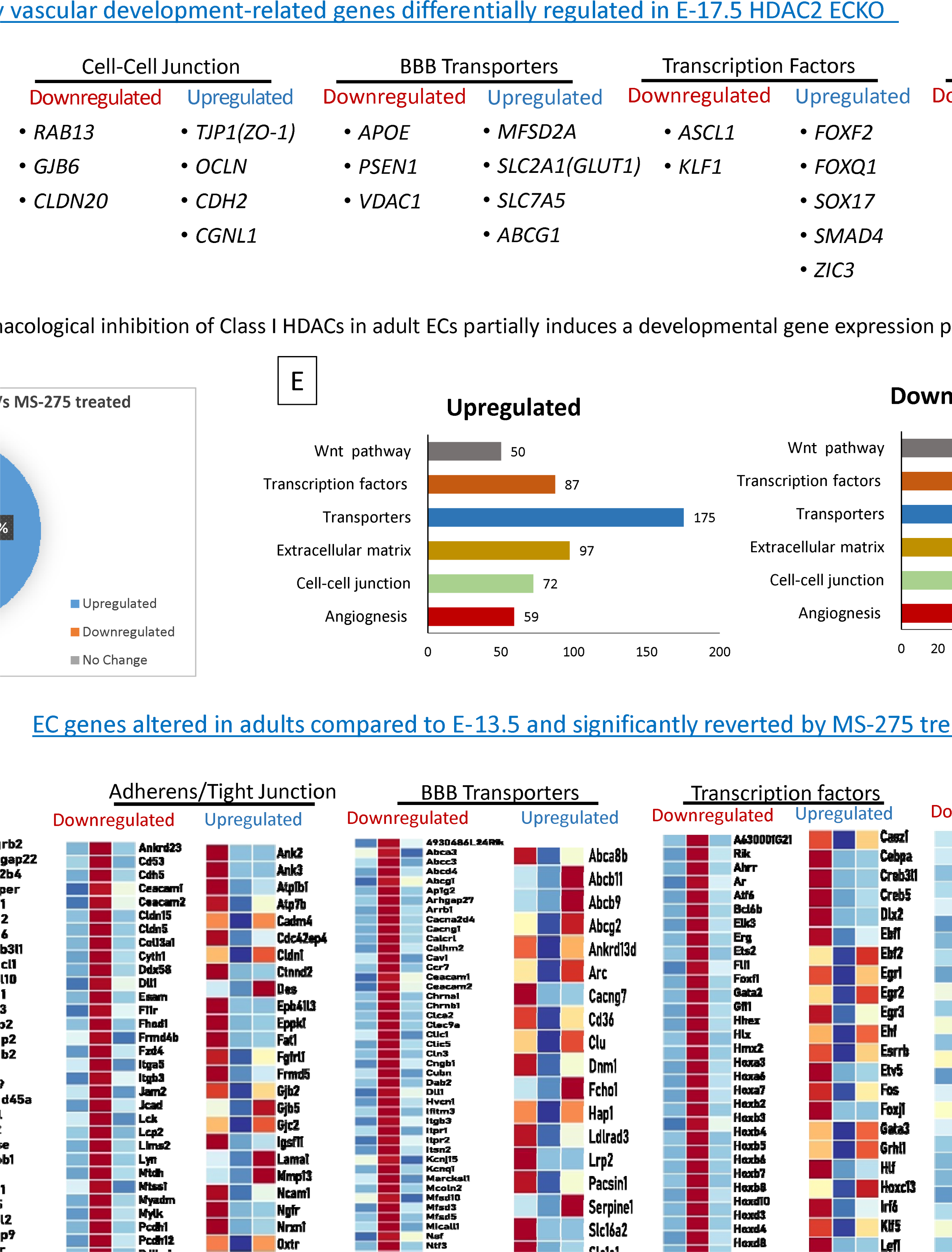
HDAC2 deletion from ECs during late embryonic development activates the angiogenesis, BBB and Wnt pathway genes. A) Tamoxifen was administered to pregnant mothers starting at E-12.5 on alternate days until E-16.5. The cortex was harvested at E-17.5, and CD31^+^endothelial cells were isolated via fluorescence-activated cell sorting (FACS). The resulting cells were processed for ultra-low-input mRNA sequencing. B) Six Key EC-regulated pathways categorize downregulated and Upregulated genes. C) Key differentially expressed genes in EC-regulated pathways. N=3 from three different mothers (*\*p* < 0.05). D) In adult ECs, treatment with MS-275 induces partial reactivation of angiogenesis and BBB formation supporting gene cohorts. Diagram depicting percentage of genes downregulated, upregulated, and shows no differential expression in MS-275 treated adult cortical ECs compared to control. E) Downregulated and upregulated genes were categorized with 6 important EC functions. F) Heat map of expression values (Z score) for differentially expressed genes (*\*p* adj < 0.05) in E13.5, adult Control and adult treated with MS-275. Five gene categories showing significant differences between E13.5 vs adult control and Adult Control vs Adult MS-275 treatment is presented. N=3 for E13.5 and adult control, N =4 for MS-275.

**Figure 8.**
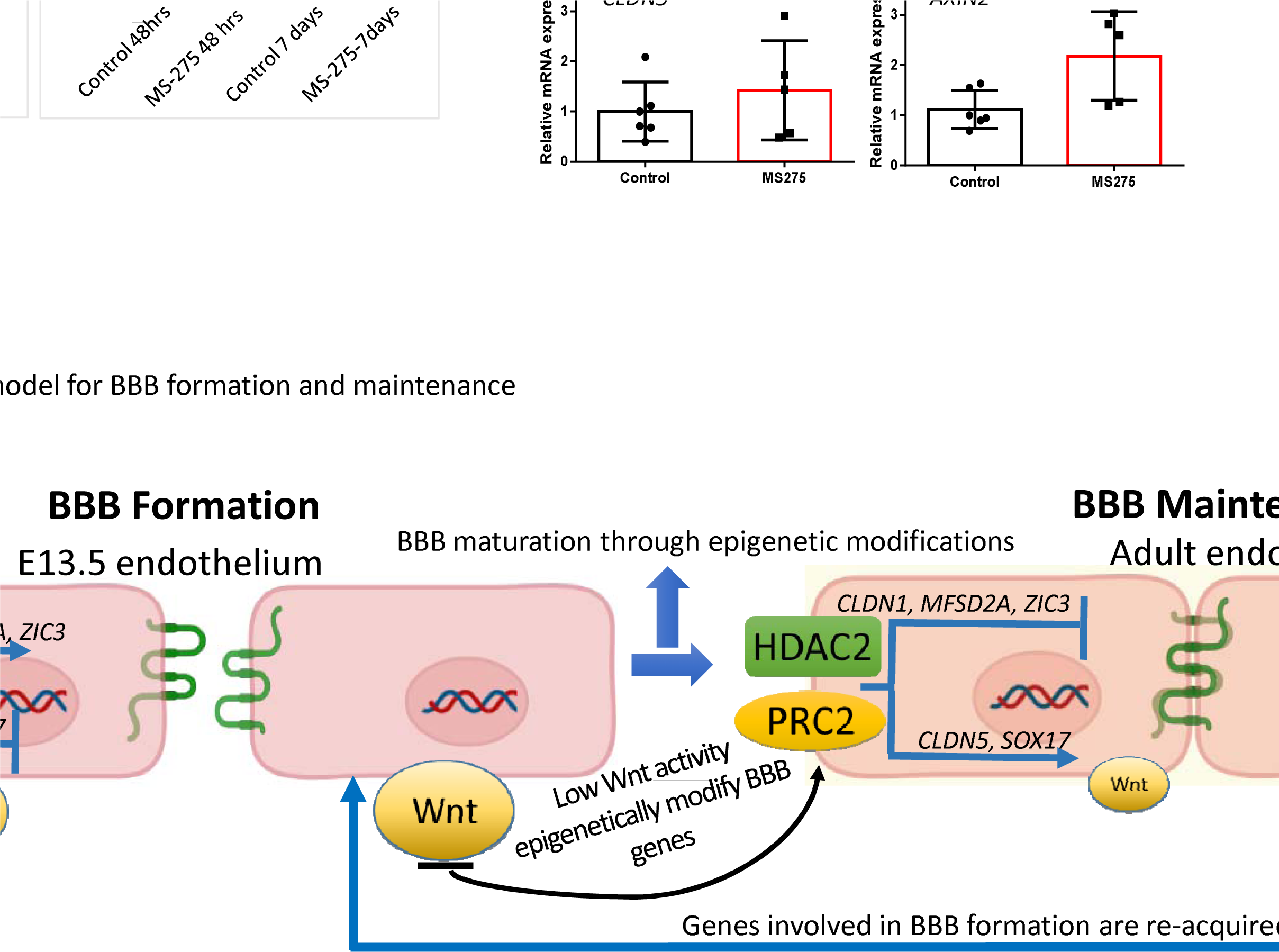
A) **Epigenetic modification induced by MS-275 is temporary** . Adult primary cortical ECs treated with MS-275 for 48 hrs. showed significant activation of *Cldn1* and downregulation of *Cldn5*. mRNA analysis on ECs after 7 days of withdrawing MS-275 showed a significant reversal of expression back to normal. *p<0.001 compared to control 48 hrs. # p<0.001 compared to MS-275 48 hrs. B) Effect of class-I HDAC inhibition on adult human cerebral arteries. Representative phase contrast image of human temporal lobe vessels in culture collected from epilepsy patients undergoing surgery. MS-275 treated human vessels showed significantly increased mRNA expression of CLDN1 and AXIN2 compared to control. *\*\*p*<0.01 and *\*p*<0.05 N=5/group. No significant difference was observed in CLDN5 expression. E) Schematic diagram illustrating the mechanisms underlying BBB formation and maintenance led by epigenetic regulators HDAC2 and PRC2. HDAC2 and PRC2 epigenetically repress EC gene cohorts that support BBB formation during development. Active Wnt signaling supports the expression of gene cohorts required for BBB formation. In contrast, a reduction in Wnt signaling recruits HDAC2 to these gene cohorts to support the formation of a functional or intact BBB. Inhibiting HDAC2 in adult ECs induces the reacquisition of gene cohorts that support BBB formation, thus representing a potential therapeutic opportunity to repair a damaged BBB.

We then performed functional classification of these genes based on key endothelial functions such as angiogenesis, cell-cell junctions, extracellular matrix, transporters, DNA-binding transcription factors, and the Wnt signaling pathway. Except for extracellular matrix function, all other categories were characterized by a significant number of upregulated genes (Fig. 7B). Key differentially regulated genes are shown in Fig. 7C. Critical angiogenesis genes, including VEGFA and ENG; tight junction proteins, including CLDN5, TJP1 (ZO-1), and OCLN; BBB transporters such as MFSD2A and GLUT1; and BBB-related transcription factors such as ZIC3, FOXF2, and SOX17 were upregulated. Further supporting the HDAC2-mediated developmental regulation of the Wnt pathway, in HDAC2 ECKO, several Wnt target genes, including AXIN2 and APCDD1, were activated in E-17.5 ECs after HDAC2 deletion. Key genes that are downregulated include APOE, PSEN1 and VEGFB.

To identify the potential of inhibiting HDAC2 in activating BBB and angiogenesis-related genes in adult ECs, and to obtain a detailed picture of CNS EC genes regulated by MS-275, a class-I HDAC inhibitor, we performed mRNA-seq analysis of MS-275-treated primary adult cortical ECs in culture and compared with the control. MS-275 treatment upregulated 48% of genes, downregulated 34% of genes, and did not alter the expression of 18% of genes.

Differentially expressed genes were grouped into six relevant categories, and all the categories showed a significant no.of differentially regulated genes (Fig-7D). Fig-7E illustrates the potential of MS-275 in regaining BBB, angiogenesis, Wnt, and transcription factors that are differentially regulated during development (E-13.5 vs Adult).

Pharmacologically Induced Epigenetic Changes Are Reversible and Partially Translate to Human Vessels We next assessed whether the gene expression changes acquired by adult ECs following MS-275 treatment were reversible. To this end, we treated adult ECs with MS-275 for 48 hours and collected ECs at 48 hours of treatment and seven days after treatment. As previously shown, *CLDN1* was significantly upregulated and *CLDN5* was downregulated considerably after 48 hours (Fig-8A). While the expression of *CLDN1* and *CLDN5* in adult ECs returned to normal in ECs collected seven days after the treatment (Fig-8A).

Finally, we examined if human vessels show similar reactivation when treated with MS-275. To this end, the human brain vessels collected from epilepsy surgery irrespective of age and gender. We tested three genes with MS-275 treatment: *CLDN1*, *CLDN5*, and *AXIN2*. We found that MS-275 treatment significantly activates the expression of *CLDN1* and *AXIN2* compared to the control, with no difference in *CLDN5* expression (Fig-8B).

## Discussion

The mechanisms that create and maintain the BBB are vitally important but poorly understood. We identified EC gene cohorts that differentially support BBB formation and maintenance, described the genetic and signaling mechanisms that establish the BBB, and presented an attractive strategy to activate gene cohorts involved in BBB formation that may promote BBB repair.

*MFSD2A* is required for BBB formation^23^, and the transcription factors *ZIC3* and *FOXF2* can induce BBB markers even in peripheral ECs^5^. Our transcriptomic analysis revealed that *CLDN1, MFSD2A* , *ZIC3*, and *FOXF2* were significantly expressed in E13.5 compared to adults, suggesting their requirement during BBB formation. Increased levels of *CLDN5*, *PECAM*, *ABCG1*, and *SOX17* in adult CNS ECs point to its significance in BBB maintenance. The results of our study partially agree with those of prior gene expression studies (S.Fig1C), but provide new concepts regarding downstream epigenetic mechanisms governing BBB gene expression. Although a high purity level is achieved in primary EC culture, we cannot exclude the possibility that transcripts from other cell populations may be identified. Even though we used similar culture conditions for both embryonic and adult cortical ECs, culture-induced changes have been reported previously ^26^ and should be considered as a varying factor when interpreting our results.

It is not known how BBB gene transcription is regulated. In CNS ECs, HDAC2 and PRC2 directly regulate the transcription of important BBB genes, including *CLDN1*, *CLDN5*, *MFSD2A*, and *ZIC3*. While HDAC2 and PRC2 commonly occupy repressed genes, we detected them in both active and repressive states at BBB genes. This result is consistent with the established dual transcriptional role of this epigenetic regulators^27–34^. Furthermore, we present evidence that HDAC2 is required to form a functional BBB and induce anti-angiogenic signals. Even though the loss of vascular integrity and lethality at E-13.5 was reported in non-inducible conditional EZH2 KO using Tie2 Cre-mouse^35^, loss of *EZH2* from E-13.5 did not significantly affect BBB permeability. These results indicate that, during the differentiation phase, HDAC2 initiates the transcriptional control, and PRC2 functions in a supporting mechanism. However, the KD of these regulators from adult CNS ECs induces a similar gene expression pattern, indicating the possibility that PRC2 facilitates the epigenetic modification during later BBB development and maintenance.

Since we did not detect H3K9me3, the repression of genes involved in BBB formation is mainly mediated through H3K27me3. On varying abundance, repressive histone mark H3K27me3 and active histone mark H3K4me3 were detected in our selected BBB genes, suggesting that repressive and activating histone methylation marks modify the promoter. Bivalent histone states can correlate with genes transcribed at low levels, suggesting these genes are poised for activation. In addition, we detected H3K9ac at the active and repressed BBB genes. Bivalent promoters can harbor H3K9ac. Thus, among the five BBB genes analyzed, each gene exhibits a different epigenetic signature, defined by histone acetylation and methylation. These results systematically demonstrate the complexity of epigenetic regulation in BBB genes and the evidence of unique epigenetic signatures. The data presented here does not entirely represent all the histone modifications on BBB genes, as other essential histone modifications, such as H3K27ac and H3K14ac, have not been examined.

The mechanism that confers low Wnt signaling in adult CNS ECs is unknown. Our results demonstrate that Wnt pathway components are still active in adult CNS ECs, yet the Wnt target genes are epigenetically inactive. Thus, in the adult CNS ECs, Wnt pathway activation permits stabilized β-catenin to enter the nucleus. Whether this nuclear β-catenin binds to Wnt target genes is unknown. Nevertheless, we demonstrated that HDAC2 and PRC2 repress Wnt target genes. Other studies have revealed that the basal or minimal Wnt pathway maintains adult BBB integrity, but its activation also prevents stroke-induced BBB damage. This also suggests the possibility of switching Wnt-regulated genes during development. Our transcriptomic analysis revealed the Wnt-regulated adult CNS EC genes associated with BBB.

It is unknown how Wnt regulates BBB genes. We demonstrated that inhibiting the active Wnt pathway at E13.5 can induce the BBB gene expression pattern associated with BBB maintenance. We demonstrate that the low Wnt pathway in E13.5 causes epigenetic modifications on BBB genes mediated by HDAC2. However, the link between Wnt and epigenetic mechanisms is unclear. Since inhibiting β-catenin induces the aforementioned results, it is likely that these effects are mediated through β-catenin. In support of this model, β-catenin GOF during BBB development prevents the maturation of BBB and inhibits the differentiation of angiogenic vessels, resulting in increased brain vascularization. Similar morphological characteristics between HDAC2 ECKO and β-catenin GOF indicate its close association.

Wnt signaling in CNS ECs is believed to require Wnt7a and Wnt7b produced by neural progenitors^36–39^. Our transcriptomic data revealed significant expression of Wnt7a in E13.5 ECs with null expression in adults. Interestingly, adult CNS ECs showed considerable activation of Wnt7a mRNA with MS-275 treatment (S.Fig7A). Moreover, β-catenin KD and GOF affect the expression of Wnt7a, and β -catenin staining after MS-275 treatment showed localization of β-catenin to the nucleus. These data indicate an innate mechanism of ECs to regulate the Wnt pathway.

A natural consequence of targeting epigenetic regulators or the Wnt pathway is that these mechanisms regulate numerous genes. Our mRNA sequencing results on embryonic and adult ECs revealed that HDAC2 inhibition or deletion could regulate EC genes associated with critical functions, including angiogenesis, barriergenesis, and Wnt signaling. Our HDAC2 ECKO phenotypic and transcriptomic data (Fig. 4A-C & Fig7A-C) reveal that the absence of HDAC2 impairs vascular and BBB maturation through the derepression of BBB, angiogenesis, and Wnt target genes (Figure 7A). The resulting increase in angiogenesis and BBB permeability in HDAC2 ECKO embryos supports our hypothesis that HDAC2-mediated epigenetic repression is essential for proper BBB and vascular development. However, our embryonic data showed no difference in CLDN1 and CLDN5 with HDAC2 deletion, which differs from the adult ECs data. This discrepancy can be attributed to developmental stage, culture-induced changes in adult ECs, pan class-I HDAC inhibitor use, heterogeneous EC population, and sequencing depth.

Epidrugs are attractive therapeutics since epigenetic changes are reversible, with the potential to reestablish function after treatment. Our results support this, as the expression of *CLDN1* and *CLDN5* returns to normal after seven days of treatment. Furthermore, our results illustrate the potential of MS-275 to reinstate the developmental characteristics in mice and partially in human adult CNS ECs.

## Acknowledgment

We thank Ralf Adams (Max Planck Institute for Molecular Biomedicine) and Ondine Cleaver (The University of Texas Southwestern) for providing the Cdh5-CreERT2 mice. This work was also supported by American Heart Association Career Development Award (18CDA34110036), NIH grant R01 to PKT (R01NS121339), and UTHealth Houston startup funds.

## Author Contributions

PKT conceived and supervised the project, designed and performed experiments, interpreted the data, and wrote the manuscript. JS and ST performed the experiments and analyzed the data under the supervision of PKT. IEM performed western blots and analyzed few RNA-seq data. HZ primarily contributed to the idea of Wnt target gene repression in adult cortical ECs. EHB provided critical comments on the manuscript and ChIP data. NT provides the human cortical vessels. AH and HL analyzed the RNA-seq data. All the authors critically commented on the manuscript.

supplementary Figure-1 Endothelial cell markers expression and validation of mRNA seq data. EC markers SLC2A1, ANGPT2 and SIPR1 expression was high in E-13.5 and adult primary cortical ECs. Adult ECs showed significant expression on CLDN5, PECAM and CDH5. These gene expressions was lower in E13.5 ECs. neuronal, astrocytes, and pericytes population was low or null in isolated ECs. B) Validation of mRNA-seq data for additional genes FOXF2 and Cldn11. C) Comparison of differentially expressed genes in E-13.5 versus adult stages with the Hupe et al. (2017) dataset, which includes genes regulated during embryonic development (E-11.5 to E-17.5). The Venn diagram shows the number of overlapping genes and highlights key genes that overlap with our data.

Supplementary Figure-2. Supplementary Figure-2. A) Western blot images and quantification showing the knockdown efficiency of HDAC2 and EZH2 sirna. *p < 0.05 vs. control N=3/4 group B) TSA treatment significantly increased the mRNA expression of ZIC3 in primary adult primary cortical ECs **p < 0.01 vs control, N=4. mRNA expression of ZIC3 and MFSD2A showed a significant increase in adult primary cortical ECs when treated with MS-275 **p < 0.01, *p < 0.05 vs control N=3. CLDN1 showed significant increase in mRNA expression in adult primary cortical ECs when treated with DZNEP. C) ChIP-qPCR showing HDAC2 and PRC2 (EED) enrichment on indicated CLDN1, CLDN5, and MFSD2A chromatin regions in E13.5 and adult cortical ECs. The ChIP signals were normalized to IgG. Data are shown as mean ± SD. ***p < 0.001, **p < 0.01, *p < 0.05 vs IgG & ###p < 0.001, ##p < 0.01, #p < 0.05 vs respective E-13.5 histone mark. N = 3-4/group.

Supplementary Figure-3. A, B,) Analysis of histone marks H3K27me3, H3K4me3, and H3K9ac in 1 KB regions of MFSD2A, ZIC3, and SOX17 genes in E13.5 and adult cortical ECs by ChIP-qPCR. MFSD2A and ZIC3 showed decreased and SOX17 showed increased expression in adults compared to E13.5. The ChIP signals were normalized to IgG. Data are shown as mean ± S.D. ***p < 0.001, **p < 0.01, *p < 0.05 vs IgG & ###p < 0.001, ##p < 0.01, #p < 0.05 vs respective E-13.5 histone mark. N= 3-4/group. D) ChIP-qPCR of H3K9me3 on indicated chromatin regions of CLDN1 and CLDN5. MFSD2A, ZIC3, and SOX17 also did not show any significant binding, and the data was not shown.

Supplementary Figure-4. A) Effect of Pharmacological inhibition of HDAC2 in WT mice. MS275(20mg/Kg) was injected IV into the pregnant mother at E13.5 and embryonic brains were harvested at E-15.5. MS- 275 treated brain showed significant growth defects in the brain with narrow cortex, shunted, and leaky vessels. Vessels were stained with IB4 and 70KDa FITC Dextran was used as a tracer. 10x images are acquired, scale bar, 100 µm. B) mRNA analysis of WT and HDAC ECKO whole brain. As a result of the tamoxifen injection, a significantly decreased expression of HDAC2 was observed in HDAC2 ECKO. BBB gene analysis revealed a significantly increased expression of CLDN1 with no difference in the expression of CLDN5, SOX17, ZIC3, and MFSD2A. **p < 0.01, vs W.T N=5/group C) Compared to W.T, EZH2 ECKO showed significantly reduced EZH2 mRNA expression. However, CLDN1 and CDN5 showed no significant difference between WT and EZH2 ECKO.

Supplementary Figure-5. a) Adult primary cortical E.C.s were treated with Wnt3a, MS-275, and MS-275

+ Wnt3a. mRNA expression of Wnt target genes AXIN2 and LEF1 showed significantly increased expression with MS-275 treatment compared to control and Wnt3a. MS-275 + Wnt3a treatment group showed significantly increased expression compared to other groups. ***p < 0.001, **p < 0.01, vs indicated conditions. N=3/condition. B) AXIN2 and LEF1 mRNA expression showed significantly increased expression in HDAC2 ECKO compared to WT **p < 0.01, vs WT N=3-5 group. C) ChIP-qPCR of H3K27me3, H3K4me3, and H3K9ac on LEF1 TSS. N d) = 3-4/group **p < 0.01, *p < 0.05 vs IgG & #p < 0.05 vs respective E-13.5 histone mark. D) Immunohistochemistry showing the expression of Wnt target Axin2 in the cortical vessels of control (DMSO +Saline), LiCl (80mg/Kg) treated and MS-275 (25mg/Kg) + LiCl treated mice. Vessels were stained with isolectin B4 (Red) and Axin2 is shown in a green channel and pointed with a white arrow. Scale bar, 10µm.

Supplementary Figure-6. A) E13.5 primary cortical ECs transfected with β-catenin siRNA showed a significant decrease in expression of AXIN2, CLDN1, ZIC3, and MFSD2A with no significant difference in CLDN5 compared to the control. **p < 0.01, *p < 0.05 Vs control N=3/group. B) Wnt target gene LEF1 showed significantly increased enrichment of HDAC2 in LF3 treated E13.5 ECS compared to control. *p < 0.05 Vs respective IgG and ##p < 0.01 Vs E13.5 control HDAC2 N=3-4/group. Repressive histone mark H3K7me3 showed significantly increased enrichment and active histone mark H3K4me3 showed significantly decreased enrichment in LF3-treated E13.5 cortical E.C.s. ***p < 0.05 Vs respective IgG and ###p < 0.001, ##p < 0.00 Vs E13.5 control H3K27me3 or H3K4me3. C) Pharmacological activation of Wnt pathway using LiCl (80mg/Kg) in pregnant mice. LiCl was injected on days as shown, and the brain was harvested at E-17.5 after the tracer was injected. White arrows show that a 70KD FITC-dextran tracer leaked outside of the vessel in the LiCl-treated embryos, and the control showed a tracer inside the vessels. Vessels were stained with isolectin B4, and the nucleus was stained with DAPI. The scale bar is 100 µm. D) mRNA analysis of AXIN2 and CLDN1 in the whole brain of WT and β-catenin GOF.

Supplementary Figure-7. A) Normalized count for Wnt7a in ECs from E13.5, adults, and adults treated with MS-275. *p < 0.05 Vs Adult and ^#^p <0.05 Vs adult. B) E13.5 primary cortical ECs transfected with β- catenin siRNA showed a significant decrease in expression of Wnt7a compared to the control. *p < 0.05 Vs control N=3/group. C) β-catenin GOF in ECs significantly increased the mRNA level of Wnt7a in the whole brain analysis. *p < 0.05 Vs WT N=3-4/group. β-catenin staining on primary adult cortical ECs treated with vehicle DMSO and MS-275(10uM). Ms-275 treatment showed nuclear localization of β- catenin into the nucleus as indicated by white arrows. 20x images are acquired. Scale bar, 20 µm.

## Notes

### Competing Interest Statement

The authors have declared no competing interest.

### Summary of Updates

To address the concern of culture-induced changes in CNS ECs and to investigate the mechanism of HDAC2-ECKO induced vascular phenotype, we have added an additional experiment in Figure 7A. We isolated CNS ECs via FACS sorting to isolate CD31+ve ECs and performed ultra-low mRNA sequencing to identify the genetic mechanisms altered in the HDAC2 ECKO phenotype. We further compared our differentially regulated genes with those identified by Hupe et al., 2017, which are regulated during development, to confirm overlapping gene sets.

