## Supplementary Figures for "Establishing And Maintaining The Blood-Brain Barrier: Epigenetic And Signaling Determinants"

### Slide 1
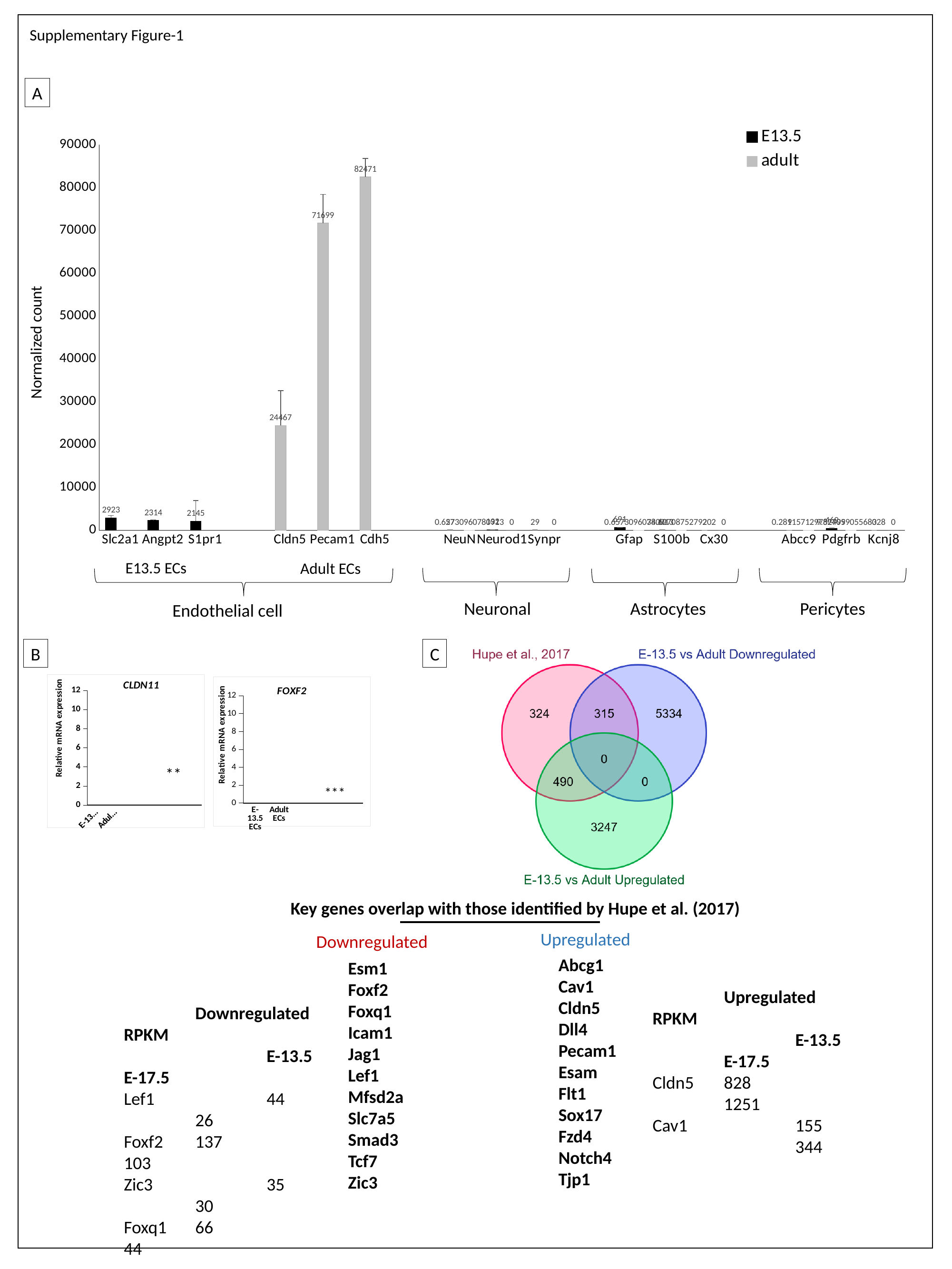

Supplementary Figure-1
A
#### Chart
| Category | E13.5 | adult |
|---|---|---|
| Slc2a1 | 2923.3248695818 | None |
| Angpt2 | 2314.43867163214 | None |
| S1pr1 | 2144.739080338683 | None |
| | None | None |
| Cldn5 | 24467.246807081687 | None |
| Pecam1 | 71699.01781363397 | None |
| Cdh5 | 82471.06324209154 | None |
| | None | None |
| NeuN | 27.43333727865159 | 0.65730960780913 |
| Neurod1 | 131.86574456327455 | 0.0 |
| Synpr | 29.07886096201503 | 0.0 |
| | None | None |
| Gfap | 691.0661731023052 | 0.65730960780913 |
| S100b | 10.326161213787033 | 34.62708752792019 |
| Cx30 | 2.167268688006347 | 0.0 |
| | None | None |
| Abcc9 | 1.2142123457906633 | 0.28915712975279534 |
| Pdgfrb | 469.14878915512 | 9.82409905568328 |
| Kcnj8 | 0.0 | 0.0 |
E13.5 ECs
Adult ECs
Neuronal
Pericytes
Astrocytes
Endothelial cell
B
C
#### Chart: CLDN11
| Category | | | |
|---|---|---|---|
| E-13.5 ECs | 1.0000000000000004 | 0.7405987244381427 | 0.25696082885309157 |
| Adult ECs | 0.33453775232165506 | 0.9437900287880787 | 0.42967907706631603 |
#### Chart: FOXF2
| Category | | | |
|---|---|---|---|
| E-13.5 ECs | 1.0000000000000002 | 0.8197504206071663 | 0.0007493981609362262 |
| Adult ECs | 0.0010880286290659753 | 0.9504349980080116 | 0.0016685575844993725 |**
***
Key genes overlap with those identified by Hupe et al. (2017)
Upregulated
Downregulated
Abcg1
Cav1
Cldn5
Dll4
Pecam1
Esam
Flt1
Sox17
Fzd4
Notch4
Tjp1
Esm1
Foxf2
Foxq1
Icam1
Jag1
Lef1
Mfsd2a
Slc7a5
Smad3
Tcf7
Zic3
	Upregulated RPKM
		E-13.5	E-17.5
Cldn5	828		1251
Cav1		155		344
	Downregulated RPKM
		E-13.5	E-17.5
Lef1		44		26
Foxf2	137		103
Zic3		35		30
Foxq1	66		44

### Slide 2
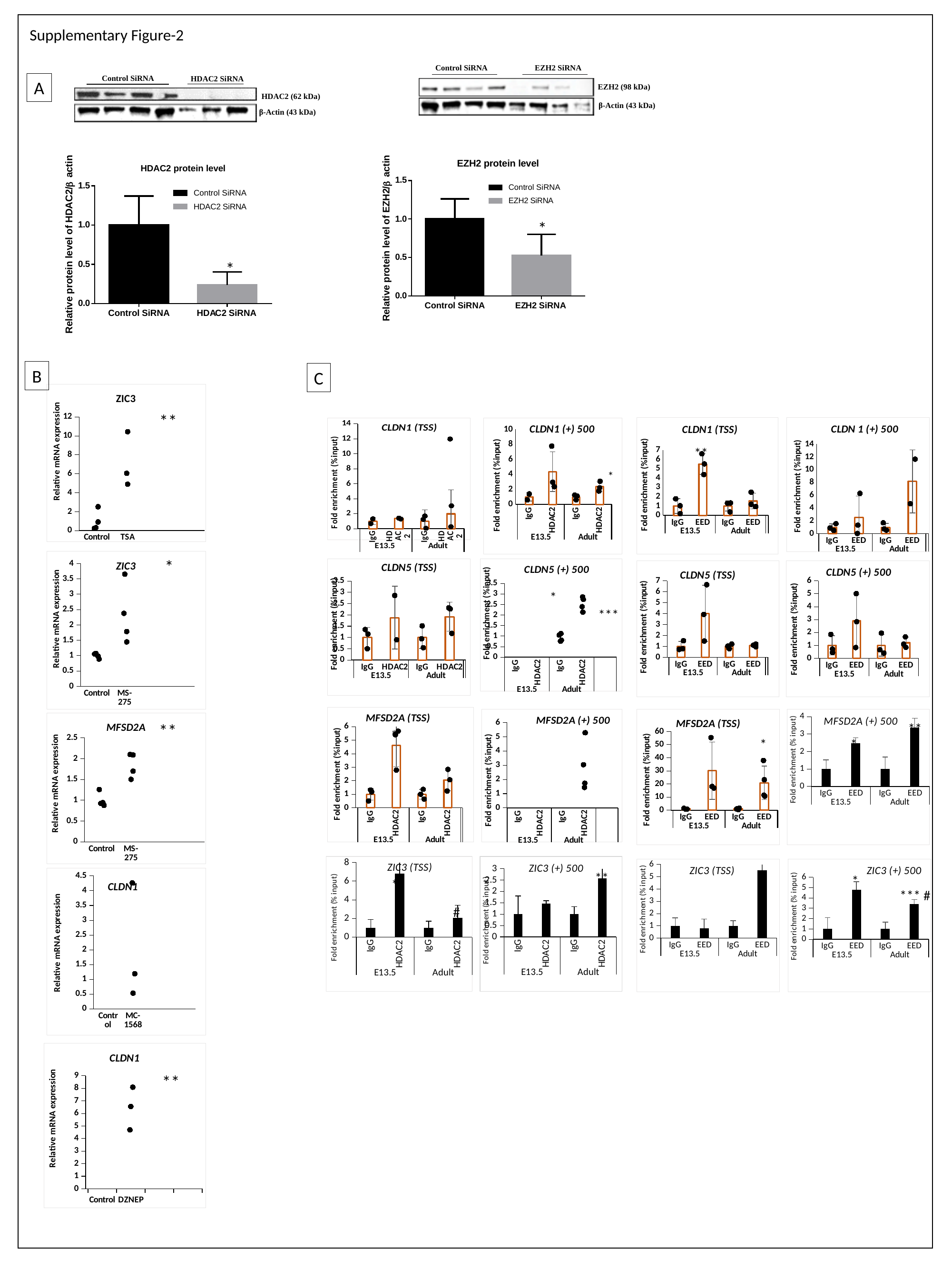

Supplementary Figure-2
Control SiRNA
EZH2 SiRNA
Control SiRNA
HDAC2 SiRNA
A
EZH2 (98 kDa)
HDAC2 (62 kDa)
β-Actin (43 kDa)
β-Actin (43 kDa)
*
*
B
C
#### Chart: ZIC3
| Category | | | |
|---|---|---|---|
| Control | 1.0000000000000013 | 0.2434175853235758 | 6.052314973050602 |
| TSA | 7.134409507254465 | 0.31830380156047955 | None |**
[unsupported chart]
[unsupported chart]
[unsupported chart]
[unsupported chart]
**
*
*
#### Chart: ZIC3
| Category | | | |
|---|---|---|---|
| Control | 1.0 | 1.0533246484747922 | 2.3783072295934606 |
| MS-275 | 2.31594108963671 | 1.070824051498651 | 1.4459605004511586 |
[unsupported chart]
[unsupported chart]
[unsupported chart]
[unsupported chart]
 *
***
[unsupported chart]
[unsupported chart]
[unsupported chart]
#### Chart: MFSD2A (+) 500
| Category | |
|---|---|
| IgG | 1.0 |
| EED | 2.4686402884748957 |
| IgG | 1.0 |
| EED | 3.376096761281326 |**
#### Chart: MFSD2A
| Category | | | |
|---|---|---|---|
| Control | 1.0 | 1.2573728342447807 | 2.100715111391481 |
| MS-275 | 1.847585973844752 | 0.9241180354698468 | 1.6993643550898323 |**
 *
*
*
#### Chart: ZIC3 (TSS)
| Category | |
|---|---|
| IgG | 1.0 |
| HDAC2 | 6.785013206630803 |
| IgG | 1.0 |
| HDAC2 | 2.0536680431099876 |
#### Chart: ZIC3 (+) 500
| Category | |
|---|---|
| IgG | 0.9999999999999999 |
| HDAC2 | 1.4657818887224805 |
| IgG | 1.0 |
| HDAC2 | 2.5768826267282976 |
#### Chart: ZIC3 (TSS)
| Category | |
|---|---|
| IgG | 0.9999999999999999 |
| EED | 0.8194151185005603 |
| IgG | 1.0 |
| EED | 5.513370258880164 |
#### Chart: ZIC3 (+) 500
| Category | |
|---|---|
| IgG | 1.0 |
| EED | 4.801967952780535 |
| IgG | 1.0 |
| EED | 3.4283423461126667 | *
**
#### Chart
| Category | | | |
|---|---|---|---|
| Control | 1.0 | 0.9690836533533466 | 4.262954630221195 |
| MC-1568 | 1.9959460734497425 | 0.31802381760248905 | 1.1904639319694474 | *
CLDN1
 *** #
#
#### Chart: CLDN1
| Category | | | |
|---|---|---|---|
| Control | 0.9999999999999991 | 0.8463413991755958 | 4.6879575721915705 |
| DZNEP | 6.443139521442927 | 0.9606000713917054 | 8.091030605430523 |**

### Slide 3
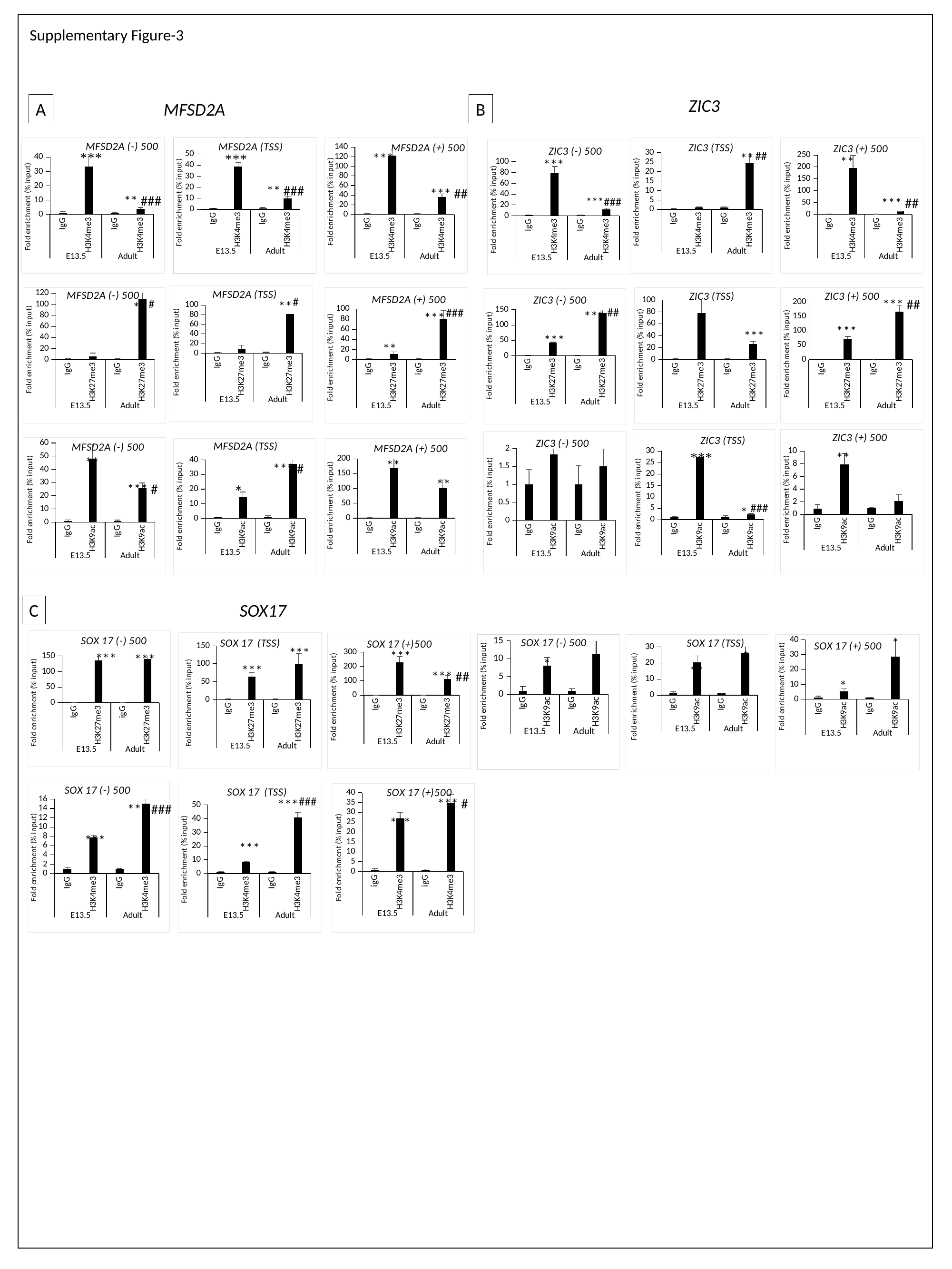

Supplementary Figure-3
ZIC3
A
MFSD2A
B
#### Chart: MFSD2A (-) 500
| Category | |
|---|---|
| IgG | 1.0 |
| H3K4me3 | 33.37818089290868 |
| IgG | 1.0 |
| H3K4me3 | 3.7697323904676194 |
#### Chart: MFSD2A (TSS)
| Category | |
|---|---|
| IgG | 1.0 |
| H3K4me3 | 38.8319805626225 |
| IgG | 1.0 |
| H3K4me3 | 9.934498812332539 |
#### Chart: ZIC3 (TSS)
| Category | |
|---|---|
| IgG | 0.2736009989820238 |
| H3K4me3 | 0.8880097155337912 |
| IgG | 1.0 |
| H3K4me3 | 24.353566535765268 |
#### Chart: ZIC3 (+) 500
| Category | |
|---|---|
| IgG | 1.0000000000000002 |
| H3K4me3 | 196.56112700862704 |
| IgG | 1.0 |
| H3K4me3 | 12.295750054607806 |
#### Chart: MFSD2A (+) 500
| Category | |
|---|---|
| IgG | 1.0000000000000002 |
| H3K4me3 | 122.95909129517963 |
| IgG | 1.0 |
| H3K4me3 | 36.3687305842337 |
#### Chart: ZIC3 (-) 500
| Category | |
|---|---|
| IgG | 1.0 |
| H3K4me3 | 78.43206826008337 |
| IgG | 1.0 |
| H3K4me3 | 12.115239891716419 |##
**
***
**
 ***
** ###
*** ##
** ###
***###
 *** ##
#### Chart: MFSD2A (TSS)
| Category | |
|---|---|
| IgG | 0.9999999999999991 |
| H3K27me3 | 8.73447178121828 |
| IgG | 1.0 |
| H3K27me3 | 80.46846696363752 |
#### Chart: ZIC3 (TSS)
| Category | |
|---|---|
| IgG | 0.9999999999999999 |
| H3K27me3 | 78.12806756959212 |
| IgG | 1.0 |
| H3K27me3 | 25.997233620423348 |
#### Chart: ZIC3 (+) 500
| Category | |
|---|---|
| IgG | 1.0 |
| H3K27me3 | 69.63544946659442 |
| IgG | 1.0 |
| H3K27me3 | 165.84693932182145 |
#### Chart: MFSD2A (-) 500
| Category | |
|---|---|
| IgG | 1.0 |
| H3K27me3 | 5.924286717799049 |
| IgG | 1.0000000000000002 |
| H3K27me3 | 109.36199626160685 |
#### Chart: MFSD2A (+) 500
| Category | |
|---|---|
| IgG | 1.0000000000000002 |
| H3K27me3 | 11.34950667291286 |
| igG | 1.0 |
| H3K27me3 | 80.68005582539372 |
#### Chart: ZIC3 (-) 500
| Category | |
|---|---|
| IgG | 1.0 |
| H3K27me3 | 41.74901139148705 |
| IgG | 1.0 |
| H3K27me3 | 138.86690074234173 |#
*** ##
#
**
**
##
###
***
***
***
***
***
**
#### Chart: ZIC3 (+) 500
| Category | |
|---|---|
| IgG | 0.8846563272592697 |
| H3K9ac | 7.885041976462369 |
| IgG | 1.0 |
| H3K9ac | 2.1331157326749763 |
#### Chart: ZIC3 (TSS)
| Category | |
|---|---|
| IgG | 1.0000000000000002 |
| H3K9ac | 27.387112001159114 |
| IgG | 1.0 |
| H3K9ac | 2.3089409104537406 |
#### Chart: ZIC3 (-) 500
| Category | |
|---|---|
| IgG | 1.0 |
| H3K9ac | 1.838596469574414 |
| IgG | 1.0 |
| H3K9ac | 1.5054759787853138 |
#### Chart: MFSD2A (-) 500
| Category | |
|---|---|
| IgG | 1.0 |
| H3K9ac | 47.90594445266469 |
| IgG | 1.0 |
| H3K9ac | 25.544049753467608 |
#### Chart: MFSD2A (TSS)
| Category | |
|---|---|
| IgG | 1.0 |
| H3K9ac | 14.49385104594277 |
| IgG | 1.0 |
| H3K9ac | 37.20346996754573 |
#### Chart: MFSD2A (+) 500
| Category | |
|---|---|
| IgG | 1.0 |
| H3K9ac | 169.10050601513242 |
| IgG | 1.0 |
| H3K9ac | 103.01368111744814 |**
**
**
*** #
**
*** #
*
 *
###
C
SOX17
 ***
 *
#### Chart: SOX 17 (-) 500
| Category | |
|---|---|
| IgG | 1.0 |
| H3K27me3 | 135.7636747667576 |
| igG | 1.0 |
| H3K27me3 | 139.90382757619804 |
#### Chart: SOX 17 (TSS)
| Category | |
|---|---|
| IgG | 1.0 |
| H3K27me3 | 64.44037317044634 |
| IgG | 1.0 |
| H3K27me3 | 98.82805900259606 |
#### Chart: SOX 17 (+)500
| Category | |
|---|---|
| IgG | 1.0 |
| H3K27me3 | 230.16101259892088 |
| IgG | 0.9999999999999998 |
| H3K27me3 | 111.60051110670405 |
#### Chart: SOX 17 (-) 500
| Category | |
|---|---|
| IgG | 1.0 |
| H3K9ac | 8.023932909228938 |
| IgG | 1.0 |
| H3K9ac | 11.172818433752772 |
#### Chart: SOX 17 (TSS)
| Category | |
|---|---|
| IgG | 1.0 |
| H3K9ac | 20.157077946508256 |
| IgG | 1.0 |
| H3K9ac | 25.919722468330374 |
#### Chart: SOX 17 (+) 500
| Category | |
|---|---|
| IgG | 1.0 |
| H3K9ac | 5.049467393016162 |
| IgG | 1.0 |
| H3K9ac | 28.621891246600427 |***
***
*
***
*
***
**
*** ##
 *
#### Chart: SOX 17 (-) 500
| Category | |
|---|---|
| IgG | 1.0000000000000002 |
| H3K4me3 | 7.754565803984884 |
| IgG | 1.0 |
| H3K4me3 | 15.060671640652181 |
#### Chart: SOX 17 (TSS)
| Category | |
|---|---|
| IgG | 1.0 |
| H3K4me3 | 8.236004601114857 |
| IgG | 1.0 |
| H3K4me3 | 41.06392018021788 |
#### Chart: SOX 17 (+)500
| Category | |
|---|---|
| igG | 1.0 |
| H3K4me3 | 26.976216510124512 |
| igG | 1.0 |
| H3K4me3 | 34.55651386526859 |###
*** #
***
*** ###
***
***
***

### Slide 4
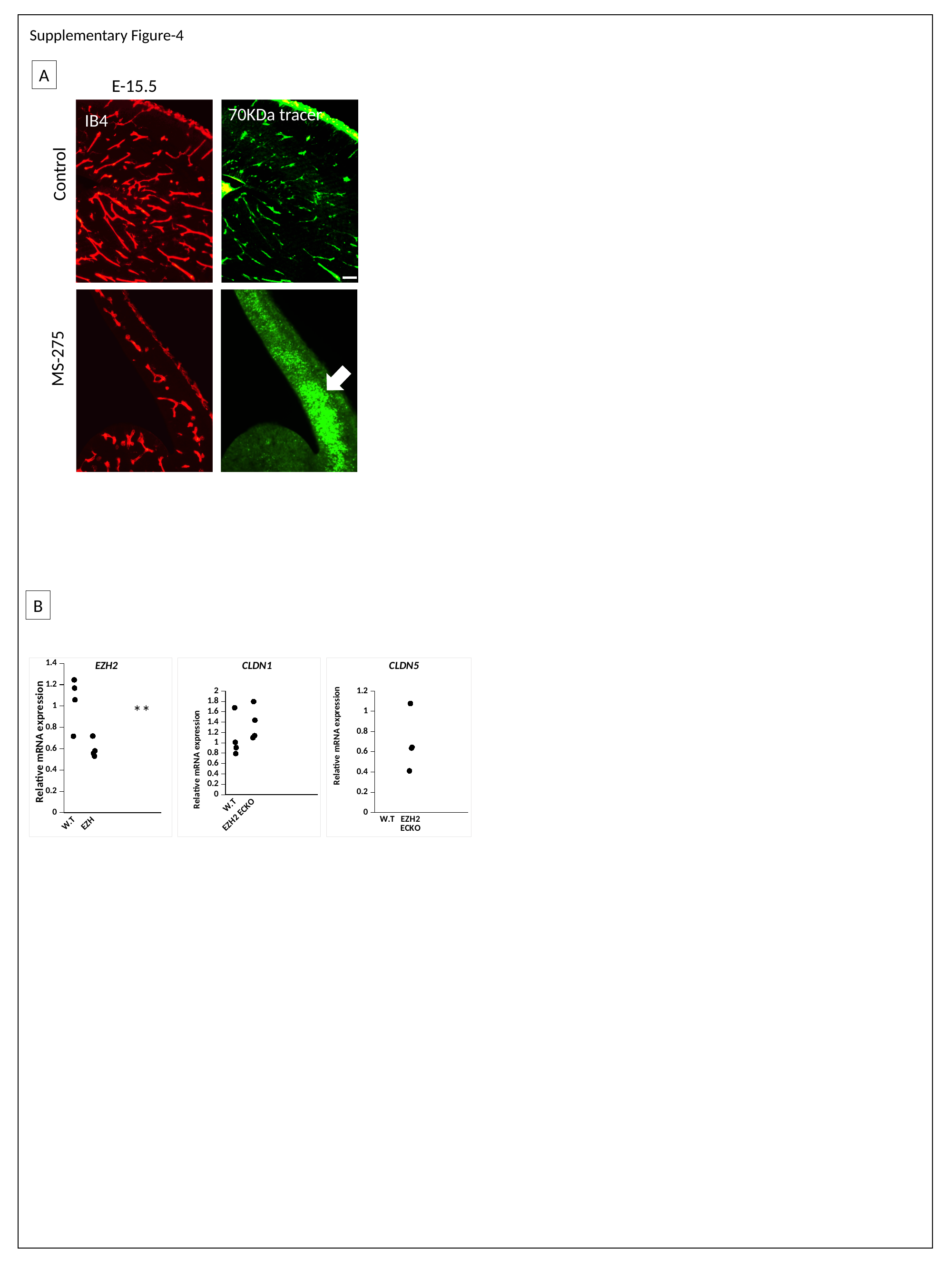

Supplementary Figure-4
A
E-15.5
70KDa tracer
IB4
Control
MS-275
B
#### Chart: EZH2
| Category | | | |
|---|---|---|---|
| W.T | 1.0 | 0.716997847394886 | 0.5555733280660379 |
| EZH2 ECKO | 0.587912265217908 | 1.2450727146118994 | 0.578776730216659 |
#### Chart: CLDN1
| Category | | | |
|---|---|---|---|
| W.T | 1.0 | 1.6778205205432655 | 1.7975249614388569 |
| EZH2 ECKO | 1.3683719783544002 | 1.0106618941330263 | 1.4370384214651293 |
#### Chart: CLDN5
| Category | | | |
|---|---|---|---|
| W.T | 1.0 | 0.9236451678693051 | 1.0768252826446603 |
| EZH2 ECKO | 0.6926266522232101 | 1.1387775968152758 | 0.6452622079690531 |**

### Slide 5
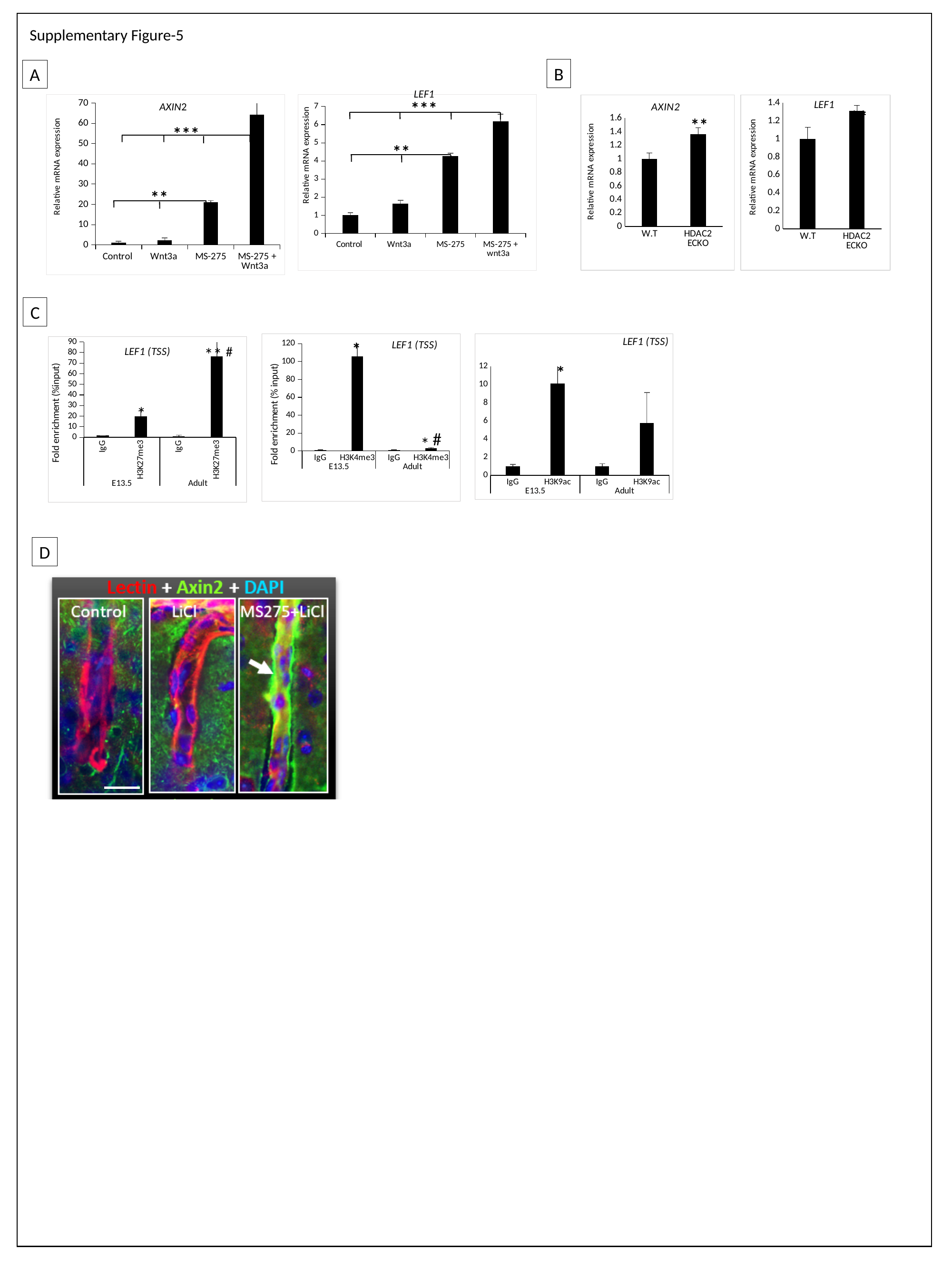

Supplementary Figure-5
B
A
LEF1
***
#### Chart
| Category | |
|---|---|
| Control | 1.0000000000000024 |
| Wnt3a | 2.352221702004077 |
| MS-275 | 20.977214988093532 |
| MS-275 + Wnt3a | 64.26828256268159 |
#### Chart
| Category | |
|---|---|
| Control | 1.000000000000001 |
| Wnt3a | 1.6493123400911156 |
| MS-275 | 4.2779781525049225 |
| MS-275 + wnt3a | 6.194454303852548 |
#### Chart: AXIN2
| Category | |
|---|---|
| W.T | 0.9999999999999998 |
| HDAC2 ECKO | 1.365143157483149 |
#### Chart: LEF1
| Category | |
|---|---|
| W.T | 0.9999999999999998 |
| HDAC2 ECKO | 1.3124316230151842 |AXIN2
**
**
***
**
**
C
#### Chart: LEF1 (TSS)
| Category | |
|---|---|
| IgG | 1.0 |
| H3K4me3 | 105.91373062461025 |
| IgG | 1.0 |
| H3K4me3 | 2.740904968882383 |
#### Chart: LEF1 (TSS)
| Category | |
|---|---|
| IgG | 1.0 |
| H3K9ac | 10.106353774806294 |
| IgG | 1.0 |
| H3K9ac | 5.783639207485776 |*
#### Chart: LEF1 (TSS)
| Category | |
|---|---|
| IgG | 1.6430653368944141 |
| H3K27me3 | 20.003004727735938 |
| IgG | 1.0000000000000002 |
| H3K27me3 | 76.35343218322949 |#
**
*
*
#
*
D

### Slide 6
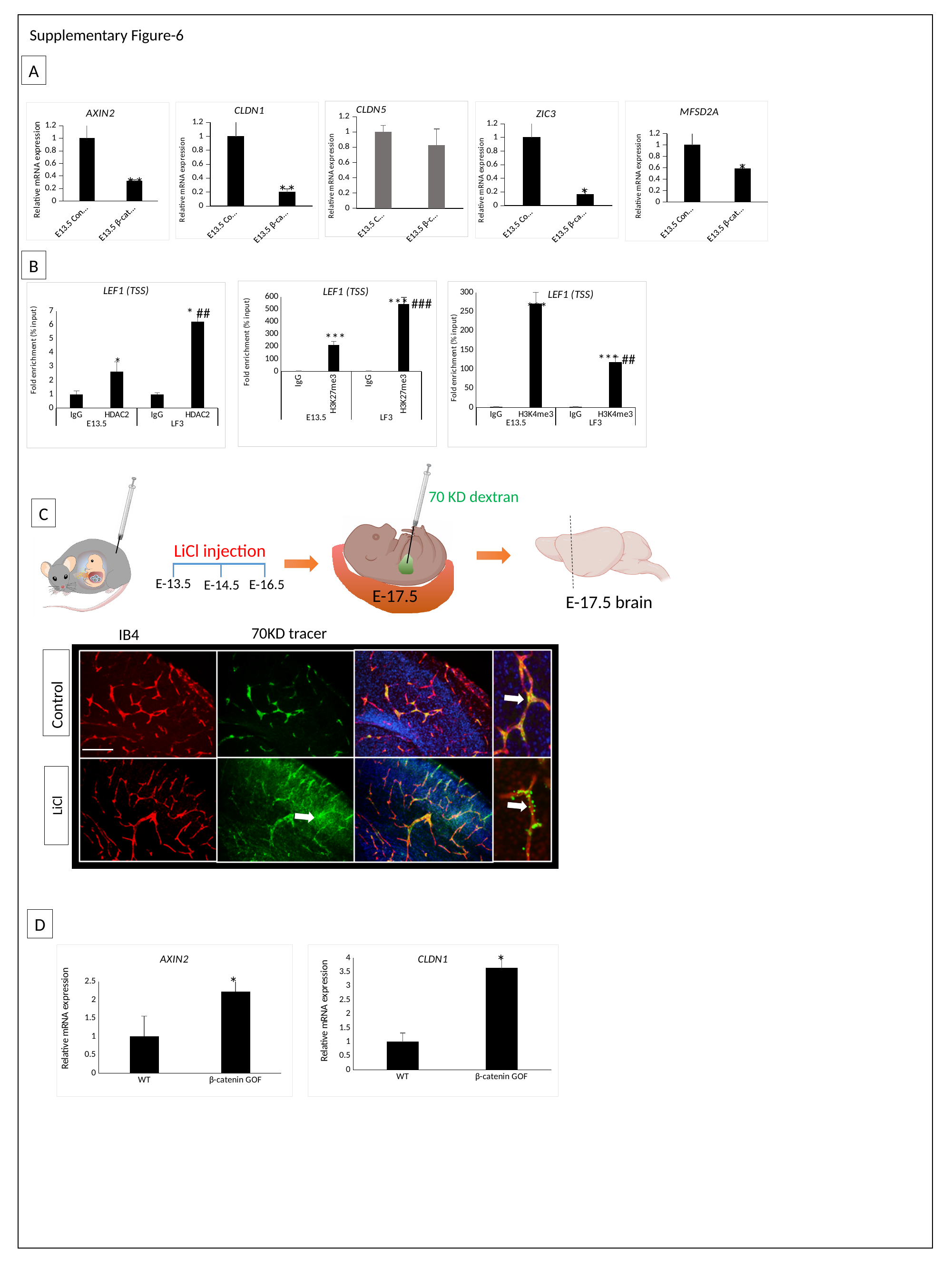

Supplementary Figure-6
A
#### Chart: CLDN5
| Category | |
|---|---|
| E13.5 Control | 1.0 |
| E13.5 β-catenin SiRNA | 0.8271298376751269 |
#### Chart: MFSD2A
| Category | |
|---|---|
| E13.5 Control | 1.0 |
| E13.5 β-catenin SiRNA | 0.585788673058655 |
#### Chart: ZIC3
| Category | |
|---|---|
| E13.5 Control | 1.0 |
| E13.5 β-catenin SiRNA | 0.16556495984210506 |
#### Chart: CLDN1
| Category | |
|---|---|
| E13.5 Control | 1.0 |
| E13.5 β-catenin SiRNA | 0.2007831438050398 |
#### Chart: AXIN2
| Category | |
|---|---|
| E13.5 Control | 1.0 |
| E13.5 β-catenin SiRNA | 0.316369528162111 |*
**
**
*
B
#### Chart: LEF1 (TSS)
| Category | |
|---|---|
| IgG | 1.0 |
| H3K27me3 | 212.66954859449947 |
| IgG | 0.75 |
| H3K27me3 | 541.481518179711 |
#### Chart: LEF1 (TSS)
| Category | |
|---|---|
| IgG | 1.0 |
| H3K4me3 | 270.33006499253776 |
| IgG | 1.0 |
| H3K4me3 | 118.38385237174818 |
#### Chart: LEF1 (TSS)
| Category | |
|---|---|
| IgG | 1.0 |
| HDAC2 | 2.6217939507300785 |
| IgG | 1.0 |
| HDAC2 | 6.24091121056011 |*** ###
***
* ##
***
*** ##
*
70 KD dextran
C
LiCl injection
E-13.5
E-16.5
E-14.5
E-17.5
E-17.5 brain
70KD tracer
IB4
Control
LiCl
D
#### Chart: AXIN2
| Category | |
|---|---|
| WT | 1.0000000000000009 |
| β-catenin GOF | 2.2224592435327346 |
#### Chart: CLDN1
| Category | |
|---|---|
| WT | 0.9999999999999999 |
| β-catenin GOF | 3.6399017970361087 |*
*

### Slide 7
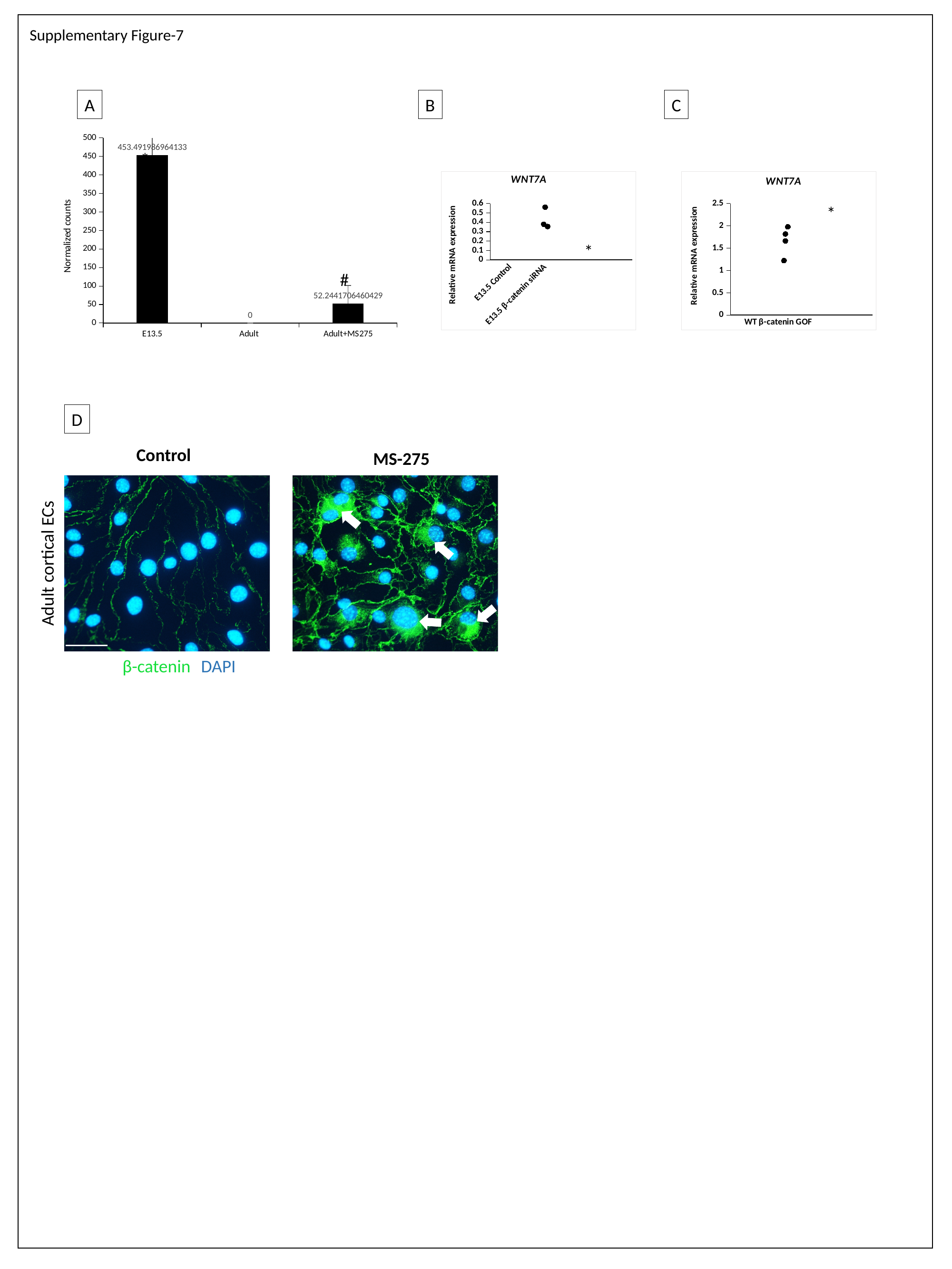

Supplementary Figure-7
A
B
C
#### Chart
| Category | |
|---|---|
| E13.5 | 453.49198696413333 |
| Adult | 0.0 |
| Adult+MS275 | 52.2441706460429 |*
#### Chart: WNT7A
| Category | | | |
|---|---|---|---|
| E13.5 Control | 1.0 | 0.8346271591217458 | 0.5621357425475336 |
| E13.5 β-catenin siRNA | 0.43238787714962496 | 0.7932906942485614 | 0.35533908048255924 |
#### Chart: WNT7A
| Category | | | |
|---|---|---|---|
| WT | 1.0000000000000002 | 1.188092694532521 | 1.8173317916040923 |
| β-catenin GOF | 1.669708533871228 | 1.0644601435286993 | 1.9782594625924987 |*
*
#
D
Control
MS-275
Adult cortical ECs
β-catenin
DAPI
